# Pre-B cell receptor acts as a selectivity switch for Galectin-1 at the pre-B cell surface

**DOI:** 10.1101/2022.02.08.479506

**Authors:** Pauline Touarin, Bastien Serrano, Audrey Courbois, Olivier Bornet, Qian Chen, Lincoln G. Scott, Stéphane J.C. Mancini, James R. Williamson, Corinne Sebban-Kreuzer, Latifa Elantak

## Abstract

Galectins are glycan binding proteins translating the sugar-encoded information of cellular glycoconjugates into many physiological activities including immunity, cell migration, and signaling. During early B lymphocytes (BL) development at the pre-B cell stage, BL express the pre-B cell receptor (pre-BCR) and are supported by mesenchymal stromal cells secreting Galectin-1 (Gal-1). Gal-1 interacts with glycosylated receptors from stromal and pre-B cell surfaces but also with the pre-BCR through a direct carbohydrate-independent contact. How this interaction might interplay with the glycan-decoding function of Gal-1 is unknown. Here, we investigated Gal-1 binding to cell surface ligands using NMR spectroscopy on native membranes. We showed that pre-BCR regulates Gal-1 binding to specifically target α2,3-sialylated receptors on pre-B cells. Upon pre-BCR interaction, dynamic changes resulted in additional contacts with α2,3-sialylated glycans converting Gal-1 from an exo- to an endo-type lectin. Remarkably, this selectivity switch is able to promote pre-B cell survival. Altogether, we shed light on a new mechanism allowing fine-tuning of Galectin specificity at the cell surfaces.

## Introduction

Glycans are essential for life and play many different fundamental roles in nearly all biological processes^1^. Their tremendous diversity is key to store chemical information at the cell surfaces, called « glycocode », which is translated into cellular responses by binding to specific proteins mainly known as lectins^2,3^. Among lectins, Galectins are a highly conserved family of fifteen known proteins (Gal-1 to Gal-15), defined by their ability to bind β-galactosides through a conserved carbohydrate recognition domain (CRD) which includes the carbohydrate binding site (CBS)^4^. These lectins, found in many cell types^5^, are well-known regulators of cell responses and mediate a wide variety of functions such as cell binding, migration, differentiation, cellular trafficking, and cell signaling^4^. Galectins are also involved in several pathological processes such as inflammatory diseases, oncogenesis, cardiovascular disorders, and host-pathogen interactions^6–10^, making Galectins attractive therapeutic targets^11–13^. All these functions, at the surface of many different cell types, in all kind of environments, are achieved through their capability to oligomerize which leads to crosslinking of specific cell-surface glycoproteins or glycolipids into lattices^14^. Given the abundance of β-galactoside-derived glycoconjugates on cell surfaces, how these lectins select a subset of ligands among many available candidates to mediate a specific cellular function remains elusive.

The concept of Galectins as functioning in the extracellular compartment only through carbohydrate interactions has been recently challenged by the identification of non-glycosylated binding partners^15–18^. Whether these interactions participate to cell glycome decoding by Galectins is unknown. The first example of a carbohydrate-independent interaction in the extracellular compartment concerns the prototype Gal-1 homodimer and the pre-B cell receptor (pre-BCR)^19^. Gal-1 is an exo-type lectin i.e. it interacts specifically with terminal units of polysaccharides^20,21^. During B cell differentiation in the bone marrow, pre-BCRs are expressed at the surface of pre-B cells, which are present in a specialized cellular niche consisting of stromal cells secreting Gal-1. At this developmental stage, Gal-1 establishes interactions with glycoconjugates and the pre-BCRs at the contact zone between pre-B and stromal cells^19,22^. This Gal-1-dependent lattice drives pre-BCRs clustering, activation, and subsequent downstream signaling which has been implicated in cell survival, proliferation, and differentiation^23^. However, the molecular mechanism underlying Gal-1 functions at the pre-B cell surface remains unknown. Our previous study revealed that Gal-1 interacts with a non-glycosylated region of the pre-BCR, the λ5 unique region (λ5-UR), that docks onto a Gal-1 hydrophobic surface behind the CBS^24^. While binding at distance from the CBS, λ5-UR induces carbohydrate affinity changes as tested on glycan arrays^15^. These data suggested that λ5-UR binding to Gal-1 modulates Gal-1 binding activity for structurally related carbohydrates, decreasing affinity for branched and linear poly-N-acetylactosamine (poly-LacNAc) and increasing affinity for sulfated or α2,3 sialylated poly-LacNAc. Whether these affinity changes represent a mechanism to regulate Gal-1 interactions and function at the pre-B cell surface is still to be demonstrated on physiological cell surface ligands embedded in the plasma membrane. To date there are no published data analyzing at the structural level Galectin binding to native cell surface ligand.

Here, we addressed the challenge of studying Gal-1 binding to pre-B and stromal cell surfaces using solution-state “on-cell” NMR spectroscopy and investigated the effect of λ5-UR interaction on Gal-1 binding to its physiological ligands. We show that λ5-UR allows Gal-1 to select specific glycosylated receptors at the pre-B cell surface. We identified α2,3-sialylated glycans as key targeted epitopes and demonstrated *in vivo* that this regulation can rescue pre-B cells from cell death. Defining the structural basis of this regulation highlighted a switch converting Gal-1 from an exo- to an endo-type lectin upon λ5-UR interaction. Altogether, our study shows that Galectin/non-glycosylated protein interactions can act as regulators of Galectin functions and may be the missing piece of the puzzle to understand how Galectins acquire their target specificity at the cell surfaces.

## Results

### NMR reveals the structural basis of Gal-1 binding to cell surface ligands

To investigate Gal-1 binding to glycoconjugates in their native membrane environment, we utilized stromal and pre-B cell lines (OP9 and Nalm6, respectively) to extract glycoconjugates-enriched membrane vesicles amenable for NMR studies. These cell lines were previously used to model the pre-B/stromal cell synapse and pre-BCR relocalization^24^. For reference, we characterized these membrane vesicle preparations by negative stain electron microscopy and observed on average 50 nm and 25 nm diameter vesicles extracted from stromal and pre-B cell lines, respectively (Supplementary Fig. 1a and 1b). NMR experiments carried out consisted in recording ^1^H,^15^N HSQC spectra on ^15^N-labeled Gal-1 alone and incubated with membrane vesicles (Fig. 1a). Upon vesicle addition, chemical shift deviations (CSDs) and line broadening were observed, indicative of complex formation with vesicle surface ligands (Fig. 1b and Supplementary Fig. 1c and 1d). To provide evidence that glycans are mediating Gal-1 binding to cell vesicles, we released complex N- and O-glycans from the cell surface by enzymatic cleavage using PNGase F and O-glycosidase. After treatment, no variation nor intensity decrease were observed on Gal-1 spectra confirming that Gal-1 interaction to cell vesicles is glycan-dependent (Fig. 1b and Supplementary Fig. 1e). These results demonstrate not only the presence of functional glycosylated ligands in the isolated membrane vesicles but also the possibility to monitor Gal-1 interactions at atomic resolution in their physiological context.

**Figure 1.**
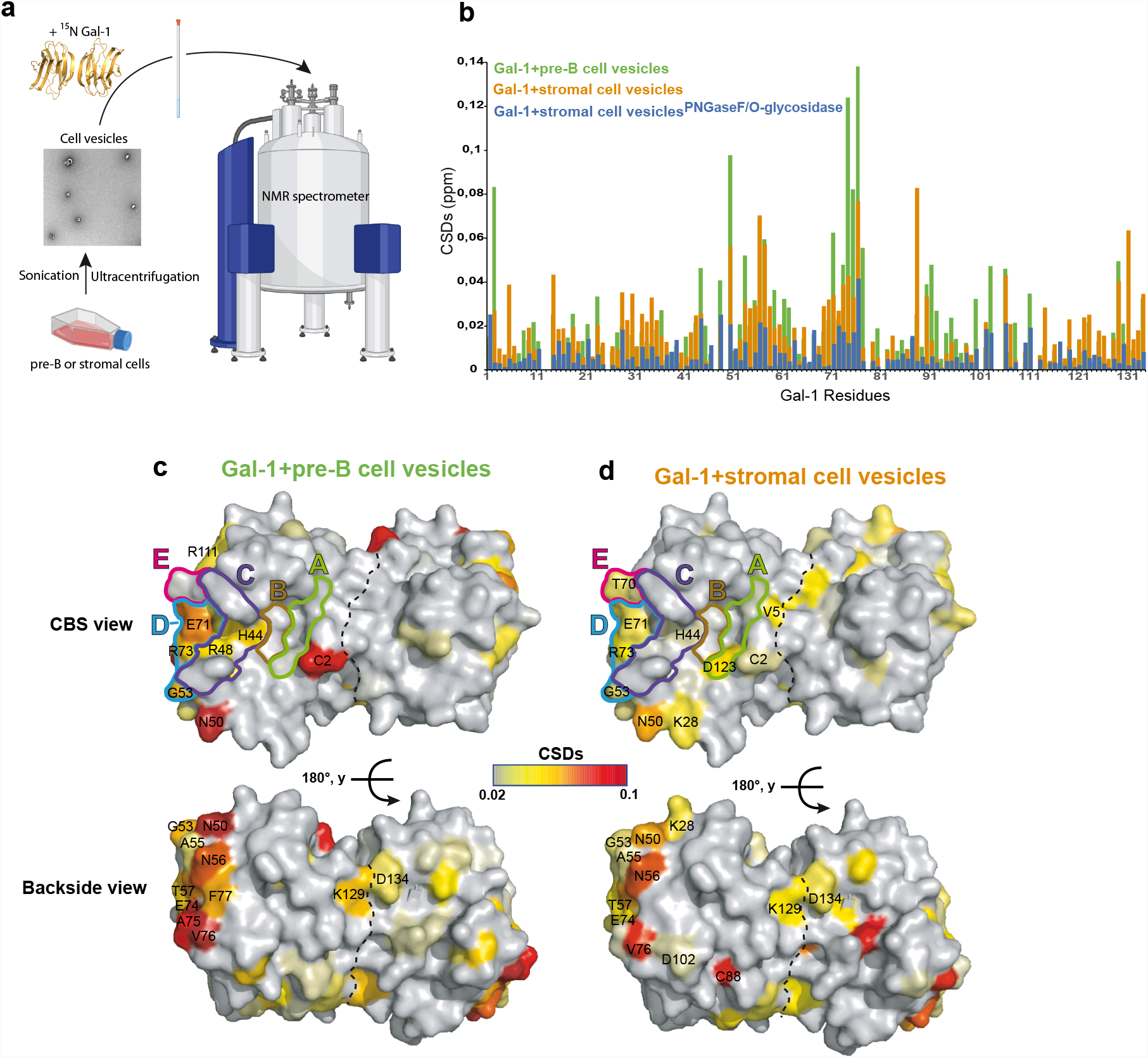
Gal-1 binding to pre-B and stromal cell surface ligands. **(a)** Strategy for on-cell NMR study of Gal-1 binding to pre-B and stromal cell vesicles. Cells are cultured as described in the Methods section, then sonicated to generate the vesicles which are collected after ultracentrifugation cycles. Vesicles are finally incubated with ^15^N-labelled Gal-1 to proceed to NMR experiment recording. **(b)** A histogram showing CSDs upon addition into ^15^N-labelled Gal-1 sample of membrane vesicles extracted from 100 million pre-B cells (green bars), 10 million stromal cells (orange bars) and 10 million PNGaseF and O-glycosidase treated stromal cells (blue bars). **(c)** and **(d)** Residues whose resonances have considerable CSDs upon **(c)** pre-B and **(d)** stromal vesicles addition are mapped onto Gal-1 homodimer surface structure (PDB: 1GZW^64^) and colored from grey (no significant CSD) to red (highest CSD) as indicated in the gradient scale. The upper panel shows the CBS view and the lower panel the backside view after a 180° rotation. CBS subsites are framed and labeled on one monomer (only one CBS is entirely visible due to the homodimer symmetry) using the following color-code: Subsite A: light green, B: brown, C: purple, D: cyan, E: magenta. Dashed lines represent the Gal-1 dimer interface.

Remarkably, Gal-1 resonances presenting CSDs are localized in the CBS (Fig. 1c and 1d). Previously, the Gal-1 CBS has been defined as containing five subsites, A to E, where subsite C is accommodating the essential β-galactose unit, the other subsites interacting more specifically with the other units of the glycan^25^. When incubated with cell vesicles, CBS subsites C and D were mainly perturbed, thus confirming Gal-1 binding to cell surface β-galactosides (Fig. 1c, 1d and Supplementary Fig. 2b and 2c). However, significant differences were observed at the residue level depending on the cell vesicles added. Remarkably, additional perturbations in CBS subsites A and E were observed in the presence of stromal cell vesicles, indicating binding to β-galactoside ligands with extensions pointing towards these subsites (Fig. 1d and Supplementary Fig. 2c). In addition to CBS perturbations, residues on the Gal-1 backside surface showed strong CSDs (Fig. 1c and 1d). These variations outside the CBS could represent an additional interaction site for extended complex cellular glycans. Several of these resonances experienced CSDs with both stromal and pre-B vesicles while some others are perturbed only with pre-B (A75, F77) or stromal (K28, C88, D102) vesicles, illustrating differences at the residue level depending on which cell vesicles were added. Finally, residues at the dimer interface showed CSDs (Fig. 1c and 1d) but also the strongest peak intensity decrease (Supplementary Fig. 2b and 2c), indicating that cell surface ligand binding to the CBS induces a long-range effect resulting in the stabilization of the dimer state. Altogether, these data show that Gal-1 binding to cell surface glycosylated ligands impact not only the CBS but also other regions of the CRD (backside and dimer interface). Ultimately, while binding to glycoconjugates containing β-galactosides, significant differences are also observed at the residue level demonstrating that pre-B and stromal cells contain different sets of Gal-1 ligands. These results are in agreement with lectin microarrays data on pre-B and stromal cells showing cell specific glycomic signatures^15^.

### λ5-UR regulates Gal-1 binding to pre-B cell ligands to target specific glycan epitopes

Within the pre-B cell niche, Gal-1 secreted by stromal cells contacts pre-B cells through binding to glycosylated receptors and to λ5-UR from the pre-BCR^19,22^. To determine whether λ5-UR can change Gal-1 binding properties to pre-B cell glycosylated ligands, we examined the effect of λ5-UR on the resulting ^1^H, ^15^N-HSQC spectra of Gal-1 in the presence of pre-B cell vesicles.

When added to Gal-1 bound to pre-B cell vesicles, λ5-UR induced CSDs in the λ5-UR binding site and within the CBS (Fig. 2a and 2b, Supplementary Fig. 3a). Remarkably, several of these resonances were already affected by pre-B vesicles interaction but, upon λ5-UR binding, they shifted in the opposite direction towards their initial Gal-1 free resonance position (Fig. 2b). Calculated CSDs for these resonances decreased until complete cancelation at a 1:3 Gal-1:λ5-UR ratio (Supplementary Fig. 3b). These observations indicate that pre-B cell ligands of Gal-1 are counter-selected in the presence of λ5-UR. Then, upon increasing λ5-UR addition, other CBS resonances located in CBS subsites B and C, and in loops surrounding the CBS started to shift are (Fig. 2b, 2c and Supplementary Fig. 3b). Consequently, the final binding pattern of Gal-1 to pre-B cell vesicles is different in the presence of λ5-UR indicating binding to new ligands (Fig. 1c vs. Fig. 2b). A control peptide, λ5-UR mutated on two essential residues for complex formation with Gal-1 (λ5-UR-L26A-W30A)^24^, has been used to verify that the effect observed is specific to λ5-UR binding to Gal-1 and not due to non-specific binding of λ5-UR to cell vesicle ligands. No CSDs were observed with the mutated peptide (Fig. 2a and Supplementary Fig. 3c). Collectively, these experiments demonstrate glycan binding changes for Gal-1 at the pre-B cell surface upon λ5-UR interaction, decreasing affinity for some glycan epitopes and enhancing interaction with others.

**Figure 2.**
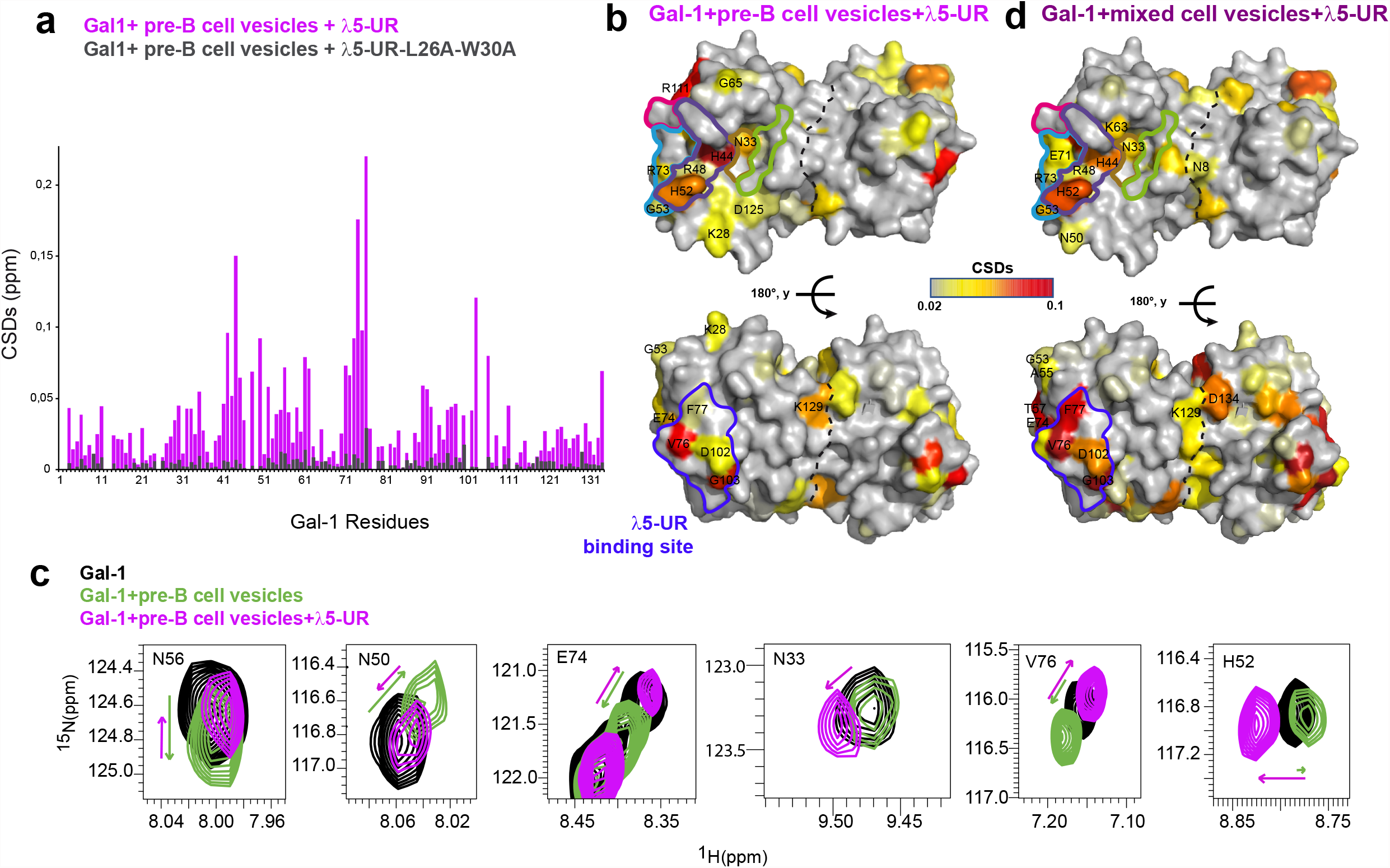
Impact of λ5-UR interaction on Gal-1 binding to pre-B cell vesicles. **(a)** A histogram showing CSDs of Gal-1 resonances upon addition of pre-B cell vesicles and λ5-UR (magenta bars), and of pre-B cell vesicles and λ5-UR-L26A-W30A (grey bars). **(b)** CSDs, calculated by comparing chemical shift resonances of Gal-1 free and Gal-1 bound pre-B cell vesicles and λ5-UR, are reported on Gal-1 surface and colored as in Fig. 1c. The λ5-UR binding site is framed in blue and labeled. **(c)** Chemical shift perturbations of selected Gal-1 resonances upon addition of pre-B cell vesicles (green) followed by λ5-UR addition (magenta). N56, N50, E74 and V76 resonances show chemical shift perturbations in the presence of pre-B cell vesicles but move back toward their initial Gal-1 free resonance position upon λ5-UR addition. N33 and H52 resonances show perturbations only upon λ5-UR interaction while being part of the CBS and not the λ5-UR binding site. **(d)** CSDs observed upon addition of mixed pre-B and stromal cell vesicles, and λ5-UR are reported on Gal-1 homodimer surface structure. As in Fig. 1, cell vesicles were extracted from 10 million and 100 million stromal and pre-B cells, respectively, which correspond for each sample to 2 mg of total protein (1:1 protein ratio).

Next, we set out to investigate what could be the λ5-UR effect on Gal-1 binding within the pre-B cell niche meaning when pre-B and stromal cell ligands are present. To mimic the pre-B cell niche, we mixed stromal and pre-B cell vesicles (Fig. 2d and Supplementary Fig. 4). Chemical shift variations observed in CBS subsites B and C were similar to the ones found when Gal-1 was titrated with λ5-UR in the presence of pre-B cell vesicles. These results indicate that, when pre-B and stromal cell ligands are present, λ5-UR regulates Gal-1 binding to target specific glycoconjugates on pre-B cells.

### Dynamic allosteric coupling mediate λ5-UR-induced Gal-1 regulation

How λ5-UR binding on Gal-1 backside is transmitted to the CBS to target specific glycoconjugates is a fundamental question. A hypothesis would be an allosteric coupling throughout Gal-1 from the λ5-UR binding site to the CBS. Allostery involves coupling of ligand binding at one site with a conformational or dynamic change at a distant site, thereby affecting binding at that site. No structural changes within the Gal-1 CBS had been observed upon binding of λ5-UR^24^, hence we investigated the role of dynamics in the regulation of Gal-1 interactions mediated by λ5-UR. The backbone dynamics of Gal-1 were analyzed using nuclear spin relaxation parameters for Gal-1 free and bound to λ5-UR (Supplementary Fig. 5). The role of fast protein motions was determined by measuring changes in the order parameter S^2^. S^2^ is a measure of the amplitude of internal motions on the picosecond-to-nanosecond timescale and can vary from S^2^ = 1, for a bond vector with no internal motion, to S^2^ = 0, for a bond vector that is rapidly sampling multiple orientations^26^. λ5-UR binding to Gal-1 caused a large number of residues to decrease their motions as evidenced by the corresponding increase in their S^2^ values (Fig. 3a). More specifically, λ5-UR binding to Gal-1 resulted in line broadening and increased rigidity (increased S^2^) for residues from the λ5-UR binding site but also for residues from the upper loop of the CBS subsite C and within subsite D and B (Fig. 3b). By contrast, increased flexibility (decreased S^2^) was observed for the lower loop of the CBS subsite C. It should be noted that chemical shift analysis showed that λ5-UR binding to Gal-1 in the presence of pre-B cell vesicles (Fig. 2b) elicited changes in chemical shift for resonances precisely located within the same regions. These findings support the idea of an allostery-based regulation of Gal-1 binding activity triggered by λ5-UR interaction in order to select specific glycan epitopes on pre-B cells.

**Figure 3.**
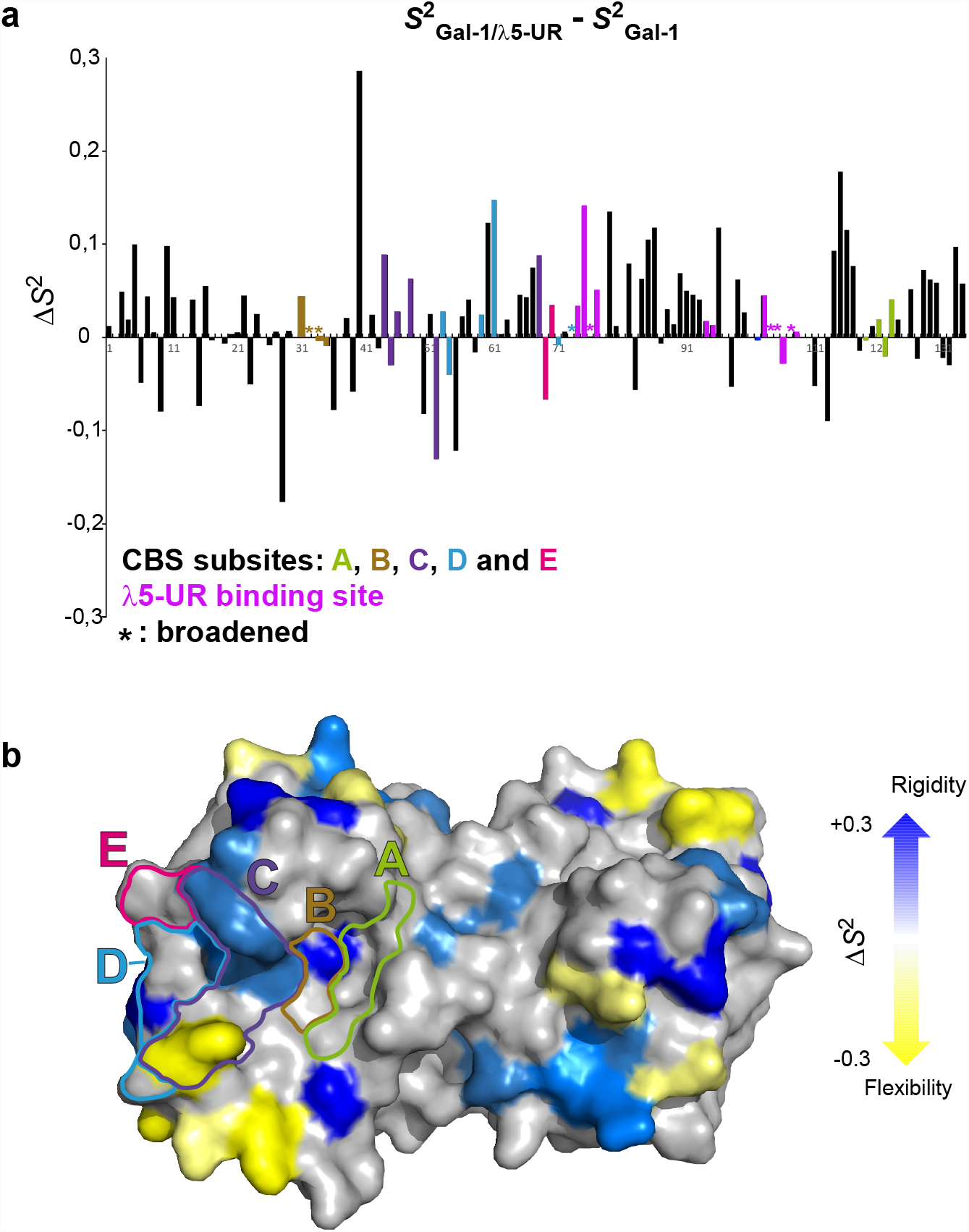
Binding of λ5-UR mediate changes in Gal-1 internal dynamics. **(a)** Changes in order parameters, Δ*S*^2^, for Gal-1 residues upon λ5-UR interaction. Δ*S*^2^ is given as *S*^2^ (after peptide binding) – *S*^2^ (before peptide binding), so positive Δ*S*^2^ values denote enhanced rigidity of the protein backbone upon peptide interaction. Residues from each CBS subsites are colored as indicated, residues involved in λ5-UR interaction are colored in magenta, and broadened peaks are indicated with asterisk. **(b)** Δ*S*^2^ values are reported on Gal-1 homodimer structure, from blue (enhanced rigidity) to yellow (enhanced flexibility).

### λ5-UR regulates binding of Gal-1 by selecting α2,3-sialylated glycan motifs found on pre-B cell surface

Our previous glycome exploration of pre-B and stromal cells highlighted a glycomic signature specific to pre-B cells corresponding to biantennary N-glycans, sulfated- and α2,3-sialylated β-galactoside glycans^15^. Importantly, the latter showed increased binding to Gal-1 on glycan arrays in the presence of λ5-UR^15^. In addition, sialylated glycans have been shown to be involved in immunological processes^27,28^. We therefore hypothesized that α2,3 sialylated glycans could be one of the specific glycosidic epitopes targeted by Gal-1 at the pre-B cell surface. To test this hypothesis on live cells, pre-B cells were incubated with the MAL II lectin which specifically recognize α2,3-sialylated glycans and is thus expected to antagonize Gal-1 binding. Remarkably, when incubated with MAL II, cells showed a strong loss in viability (Fig. 4a) indicating that interaction of MAL II with the broad range of α2,3-sialylated receptors at the pre-B cell surface triggers a signaling pathway leading to cell death (Fig. 4b). Neither Gal-1 nor λ5-UR reversed the induction of cell death by MAL II, thus illustrating their individual inability to perturb MAL II interactions with α2,3-sialylated receptors (Fig. 4a and 4c). However, when Gal-1 is combined to λ5-UR, the MAL II-induced cell death was significantly inhibited (Fig. 4a). These observations demonstrate that λ5-UR binding to Gal-1 provokes increased affinity for α2,3-sialylated receptors containing β-galactosides, perturbing MAL II network interactions and precluding cell death signaling (Fig. 4d). Of note, in the absence of MAL-II, neither Gal-1, λ5-UR or a combination of both are able to induce strong pre-B cell death as seen with MAL II (Fig. 4a). Altogether, these results demonstrate that λ5-UR mediates Gal-1 specific targeting of ligands containing α2,3-sialyl moieties at the pre-B cell surface and can promote pre-B cell survival. The structural basis of such ligand binding selection remains to be unraveled.

**Figure 4.**
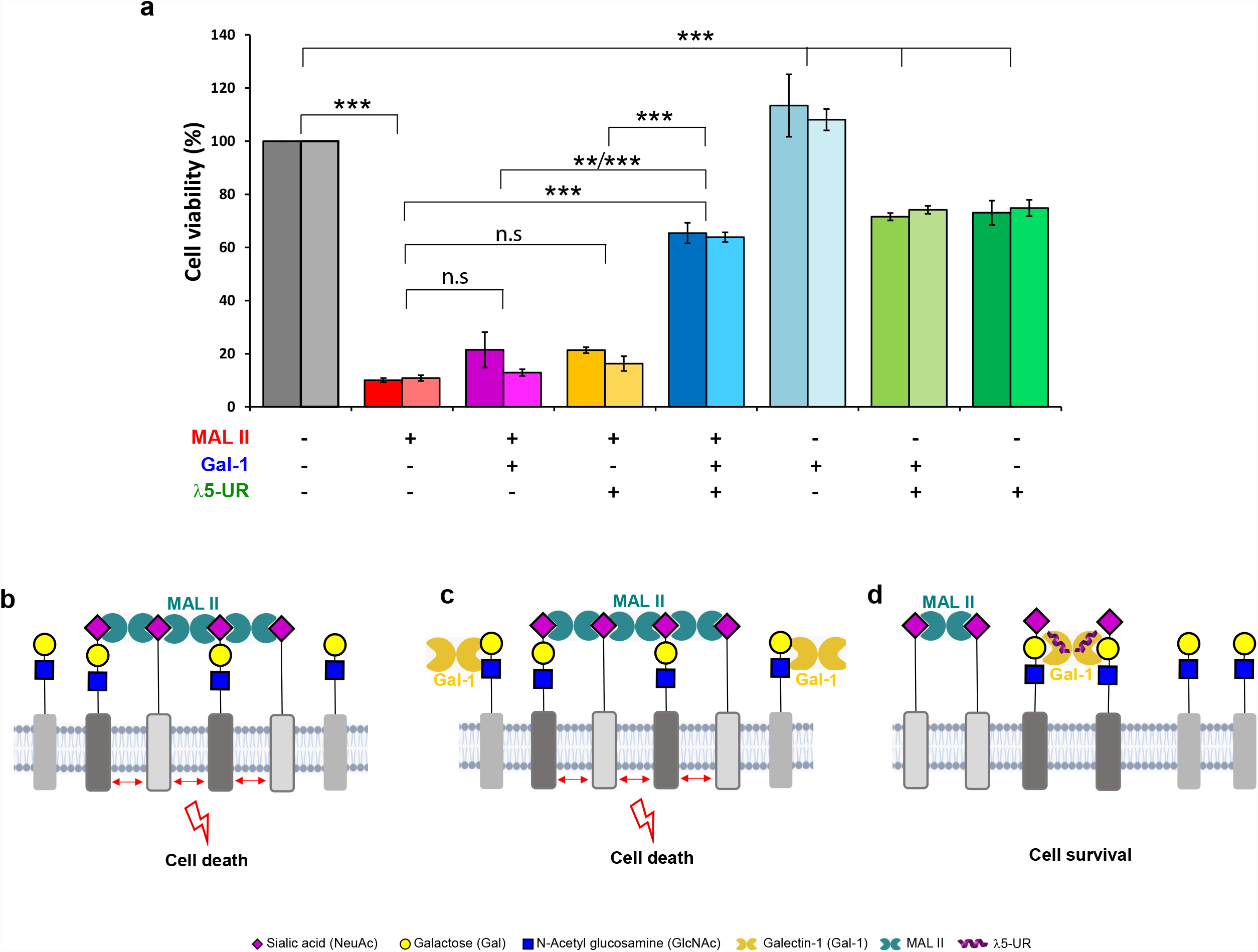
Gal-1 specifically targets α2,3 sialylated glycans at the pre-B cell surface when bound to λ5-UR. **(a)** Histogram plot showing Pre-B cell viability after incubation with MAL II, Gal-1 and λ5-UR as indicated below the histogram. For each condition two bars are shown, the left bar corresponds to pre-B cell viability evaluated using CellTiter-Glo Luminescent Cell Viability Assay (Promega) and the right bar using Trypan blue exclusion test. Means and standard errors from three independent experiments are presented. Significant differences are represented as *p<0.05, **p<0.01 and ***p<0.001. n.s indicates no significant differences. **(b)** Schematic illustrating MAL II induced pre-B cell death through clustering of α2,3 sialylated receptors. **(c)** The presence of Gal-1 does not perturb MAL II interactions and therefore pre-B cell death. **(d)** λ5-UR binding to Gal-1 increases Gal-1 binding to α2,3 sialylated receptors containing β-galactosides, thus perturbing MAL II interactions and allowing Pre-B cell survival. Glycans are represented using the symbol nomenclature as indicated^65^.

### λ5-UR induces increased Gal-1 contacts with α2,3 sialylated di-LacNAc, converting Gal-1 from exo- to endo-type lectin

To further investigate at atomic resolution the effect of λ5-UR on the Gal-1/α2,3 sialylated glycan interaction, we synthesized using a chemo-enzymatic based methodology the α2,3 sialyl di-LacNAc pentasaccharide called hereafter SdiLN (Supplementary Fig. 6a). Since galactose moieties are essential to Gal-1 binding, we also synthesized the pentasaccharide fully ^13^C-labeled on the galactose moieties (Supplementary Fig. 6). First, we tested its interaction with Gal-1 using saturation transfer difference (STD) NMR spectroscopy. This technique is based on selective irradiation of protein protons and subsequent detection of magnetization transfer to small ligands^29,30^. In the presence of Gal-1, magnetization transfer was observed indicating intermolecular contacts with the pentasaccharide (Fig. 5a). The STD signal mainly arises from interaction with the terminal galactose (Gal_A_) as expected for an exo-type lectin (Fig. 5a and Supplementary Fig. 7a). In addition, ^1^H,^13^C-HSQC spectra recorded before and after addition of Gal-1 to the ^13^C-labeled SdiLN showed severe peak loss for Gal_A_ which is therefore the core binding unit to Gal-1 (Fig. 5b and 5c). These data are consistent with the Gal-1 classification as an exo-type lectin^31,32^. Remarkably, while only galactose moieties were ^13^C-labeled, peaks corresponding to resonances from the terminal N-acetyl glucosamine (GlcNAc) and sialic acid (NeuAc) were visible on the HSQC spectrum in the presence of Gal-1 (Fig. 5b). This observation suggests that binding to the protein changes the dynamic of these groups resulting in a higher flexibility which, combined to the HSQC experiment sensitivity, make them visible on the spectrum despite the weak ^13^C natural abundance. The reverse experiment, where Gal-1 is ^15^N-labeled and the SdiLN is unlabeled has also been performed and showed CSDs and decreased peak intensities mainly for Gal-1 resonances located in CBS subsite C where the core binding galactose Gal_A_ of the SdiLN is likely interacting (Fig. 5d and Supplementary Fig. 8). Surprisingly, in the presence of λ5-UR, STD signal contributions became equivalent for Gal_A_ and Gal_B_ demonstrating that Gal-1 interaction is not exclusive to the terminal galactose when bound to λ5-UR (Fig. 5e and Supplementary Fig. 7b). As well, ^1^H,^13^C-HSQC spectrum of SdiLN showed loss of Gal_A_ and Gal_B_ resonances upon addition of λ5-UR confirming interaction with both terminal and internal galactose (Fig. 5f and 5g). Moreover, NeuAc and GlcNAc resonances diseappeared indicating increased contacts with these glycan moieties. On the Gal-1 side, addition of λ5-UR to the Gal-1/SdiLN complex not only intensified the initial perturbations but also propagated the effect towards CBS subsites A to E illustrating increased intermolecular contacts throughout the CBS with the pentasaccharide (Fig. 5h and Supplementary Fig. 8). λ5-UR alone (without the SdiLN) is not able to prompt such Gal-1 signal intensity decrease which is consistent with the binding of a 24 amino acid peptide in fast exchange regime^13^. Thus, λ5-UR interaction modifies Gal-1 binding to α2,3 sialylated glycan by allowing additional contacts with the pentasaccharide including with the internal galactose unit. While recent studies have elegantly rubber-stamped the selectivity of Gal-1 towards terminal galactose^21^, here we establish that λ5-UR binding to Gal-1 induces a selectivity switch converting Gal-1 from an exclusive exo-type lectin to an endo-type lectin interacting with both terminal and internal galactose units (Fig. 5i). This switch is therefore at the basis of Gal-1 target specificity at the pre-B cell surface.

**Figure 5:**
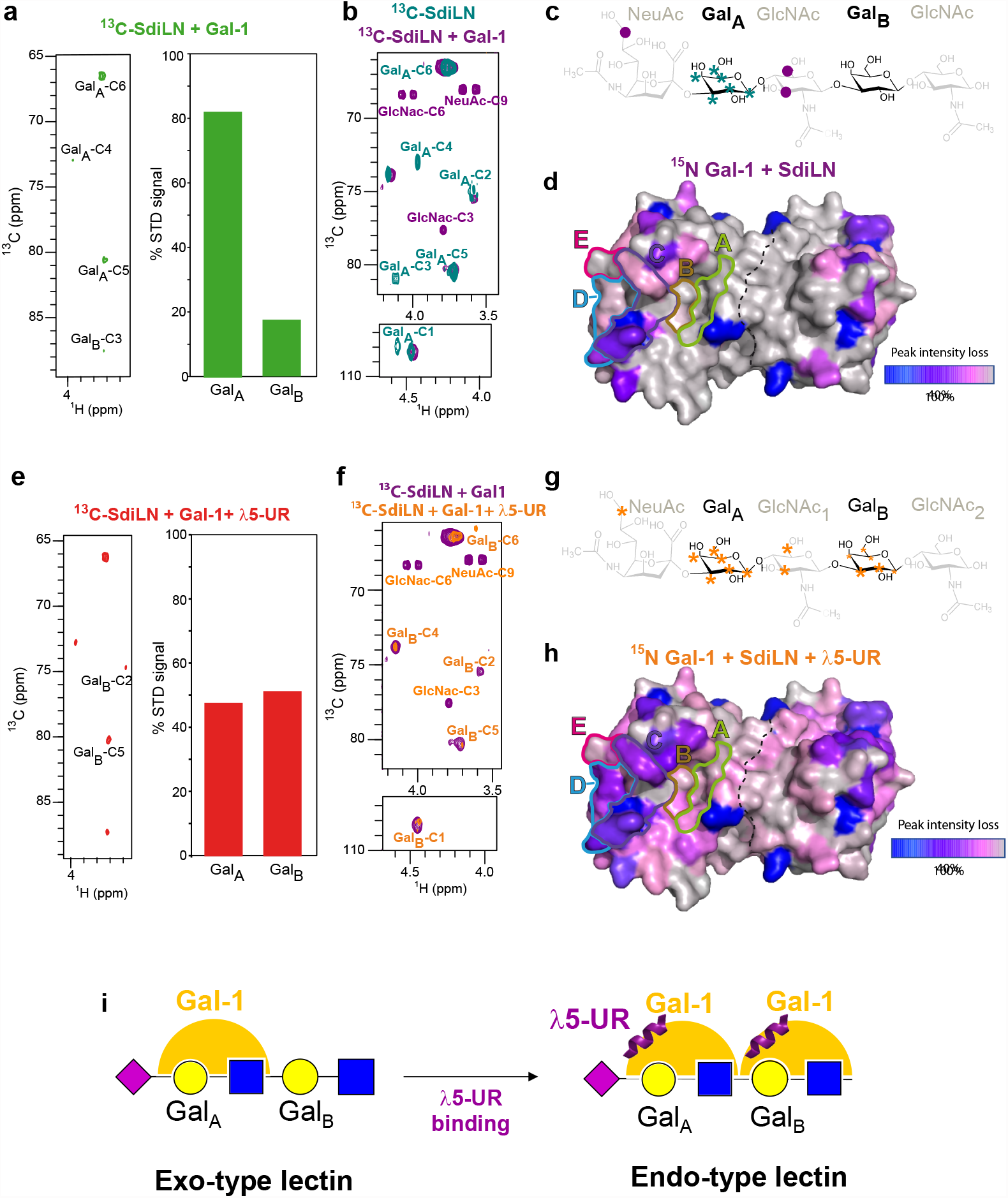
λ5-UR binding to Gal-1 induces enhanced and additional intermolecular contacts with α2,3 sialyl di-LacNAc (SdiLN). **(a)** ^1^H,^13^C STD-HSQC spectrum (left) of 1mM SdiLN ^13^C-labeled on galactose moieties in the presence of 10µM Gal-1. Contribution to Gal-1 binding for each galactose as a percentage of the total STD signal observed is shown (right). **(b)** Overlay of ^1^H,^13^C HSQC of labeled SdiLN (teal) and SdiLN in the presence of equimolar amount of Gal-1 (purple). Peaks disappearing are labeled (teal labels). Peaks appearing are also labeled (purple labels). **(c)** Variations are reported on the SdiLN chemical structure with unlabeled moieties greyed out. Teal stars indicate SdiLN C-H groups for which resonances disappear upon Gal-1 binding. Purple circles indicate SdiLN groups for which resonances appear upon Gal-1 binding. **(d)** Peak intensity loss observed on ^1^H,^15^N HSQC spectrum of Gal-1 after addition of SdiLN is reported on Gal-1 homodimer surface structure. A color gradient has been used to indicate the percentage of intensity loss as indicated on the gradient bar. CBS subsites are framed and labeled A to E. **(e) to (h)** Same as in **(a)** to **(d)** but after addition of λ5-UR. **(e)** ^1^H,^13^C STD-HSQC spectrum (left) of 1mM SdiLN ^13^C-labeled on galactose moieties in the presence of 10µM Gal-1 and 20µM λ5-UR. Contribution to Gal-1 binding for each galactose as a percentage of the total STD signal observed is shown (right). **(f)** Overlay of ^1^H,^13^C HSQC of labeled SdiLN bound to Gal-1 (purple) and after addition of λ5-UR (orange). Peaks disappearing or showing decreased intensity are labeled and shown in orange. **(g)** Variations are reported on the SdiLN chemical structure. Large orange stars indicate SdiLN C-H groups for which resonances disappear upon Gal-1 and λ5-UR binding. Small orange stars indicate groups showing decreased peak intensity. **(h)** Peak intensity loss observed on ^1^H,^15^N HSQC spectrum of Gal-1 bound to λ5-UR after addition of SdiLN is reported on Gal-1 surface structure. **(i)** Schematic illustrating the Gal-1 selectivity change upon λ5-UR interaction. Without λ5-UR, Gal-1 acts as an exo-type lectin, binding to the terminal galactose. Upon λ5-UR binding, Gal-1 is able to interact with the internal galactose and thus Gal-1 is converted into an endo-type lectin.

## Discussion

Over the past decades, many cellular activities have been ascribed to Galectins^4^. Such a wide range of functions in so many cell types and environments have conducted many laboratories to develop methodologies to understand the varied and intricate roles played by these ubiquitous lectins. While many of these experiments developed provide evidence and enable visualization for Galectin–glycoprotein lattices on cell surfaces^33,34^, further studies were needed to understand the Galectins structural features regulating lattice assembly/disassembly at the cellular level. Here, we directly examined for the first time the structural properties of Gal-1 binding to native membranes by combining solution-state NMR spectroscopy to the preparation of pre-B and stromal membrane vesicles. Importantly, we could demonstrate that Gal-1 binding to its physiological cellular ligands involves long-range effects throughout the CRD beyond the canonical CBS limits. While multivalency of Gal-1 and its glycoconjugate ligands is essential for the formation of supramolecular lattices^35^, the additional interaction surface revealed for extended complex sugar (Fig. 1c and 1d) as well as the conformational exchange observed (Supplementary Fig. 2b and 2c) for residues at the dimer interface could be a key mechanism allowing increased overall stability and biological functionality of the assembled lattice on the cell surface.

Remarkably, our on-cell NMR approach allowed us to address how Gal-1 is able to decode the glycome of pre-B cells. Indeed, given the abundance of β-galactosides on the cell in the form of glycoproteins and surface glycolipids, one would expect that Galectin/glycoconjugates lattice translates into a large heterogeneous complex on the surface of cells. But conversely, this lectin is able to cross-link a single species of glycoproteins to form a uniform lattice^34,36,37^. Here, we demonstrate that Gal-1 is able to achieve target specificity at the pre-B cell surface through a direct interaction with a non-glycosylated protein domain of the pre-BCR, λ5-UR. This interaction allows Gal-1 to interact with α2,3 sialylated glycoconjugates. Sialic acids are ubiquitously and abundantly found on the surface of all human cells as the terminating sugar in glycolipids (gangliosides) and glycoproteins (complex N-glycans and mucin type O-linked glycans). Moreover, sialic acids have been shown to play essential roles in immunological processes and in particular during B cell receptor (BCR) signaling^28,35,38–40^. In this context, the α2,6 sialylated CD22 receptor, which is a lectin recognizing α2,6 sialylated glycoreceptors, is recruited close to the BCR and inhibits its downstream signaling (Fig. 6a). The objective of this control being to prevent inadvertent activation to weak signals that could be considered a form of self-recognition and/or unwanted B cell responses under the appropriate circumstances^40^. Our results suggest that Gal-1 bound to λ5-UR would help recruitment of α2,3 sialylated receptors at the pre-B synapse. In addition, although our experimental setup using MAL II lectin is not a physiological condition, we showed that clustering of a broad range of α2,3 sialylated receptors lead to pre-B cell death and that Gal-1/ λ5-UR complex is able to disrupt the clustered sialylated receptors thus promoting pre-B cell survival (Fig. 3). Similarly to the CD22 dependent regulation of BCR signaling, it is tempting to propose that MAL II in our experiments would mimic a physiological receptor which would trigger specific cellular responses (Fig. 6b). The presence of Gal-1 bound to the pre-BCR would disrupt this clustering by recruiting specific α2,3 sialylated receptors at the pre-B synapse to activate proper signaling (Fig. 6b). Indeed, restricted receptor segregation into membrane microdomains would result in a homogenous platform suitable for efficient pre-BCR signaling leading to proliferation and survival. In line with this model of pre-BCR activation, it has been previously observed that Gal-1 binding resulted in the formation of large highly immobile pre-BCR aggregates at the pre-B cell surface^41^. Moreover, the clustering of specific glycoproteins and/or glycolipids could generate mechanical stress which translates into membrane curvature necessary for pre-BCR internalization following cell signaling. This model is strongly supported by previous evidence showing that another Galectin, the chimeric Galectin-3 (Gal-3), is able to mediate endocytic invaginations by helping the clustering of cargo proteins and glycosphingolipids^39,40^. Overall, our study reveals that the Gal-1/pre-BCR interaction is part of the glycome decoding mechanism on the surface of pre-B cells and would aim to re-orchestrate receptors localization to ensure cell signaling regulation and proper B cell development.

**Figure 6.**
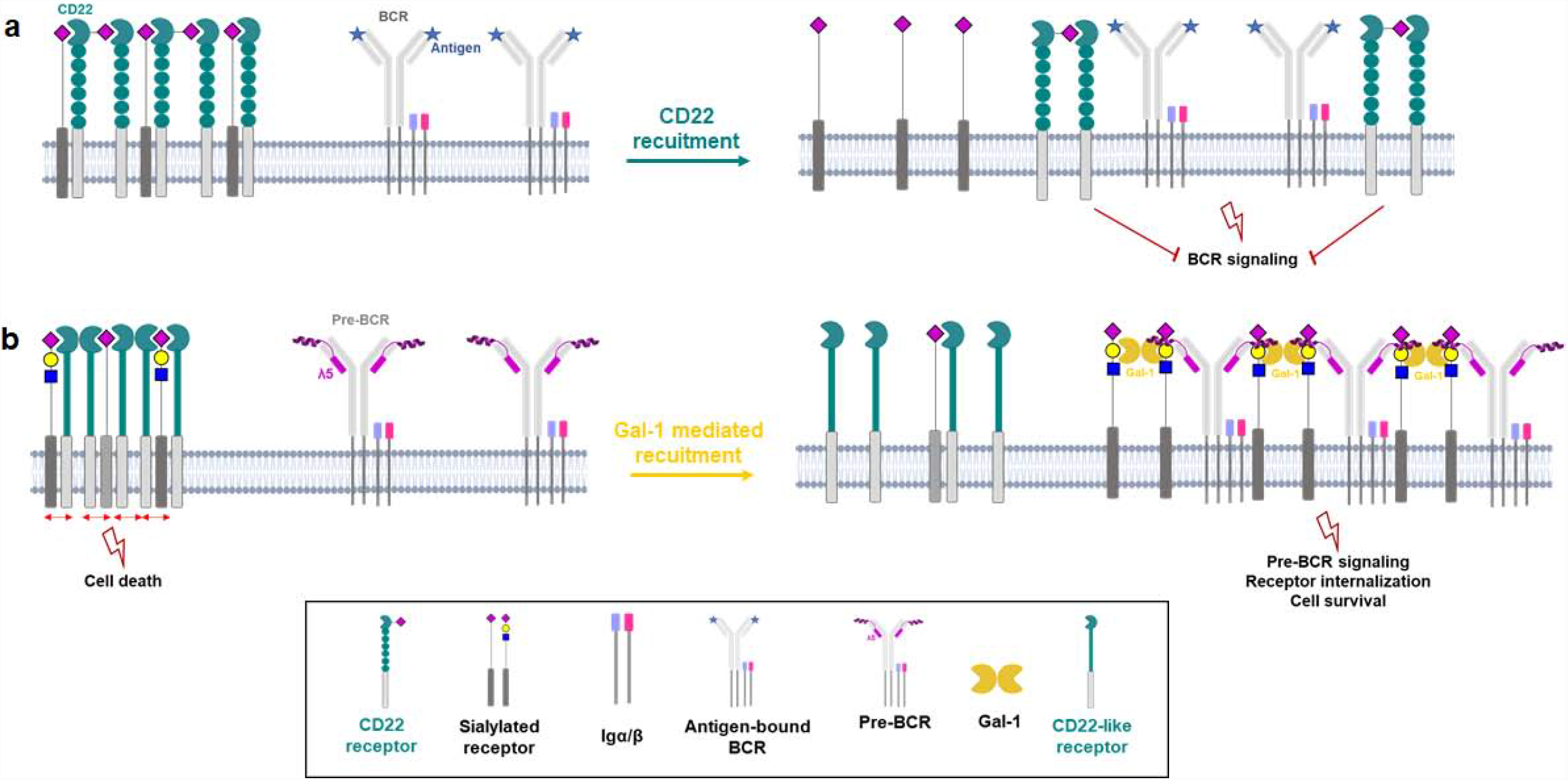
Model of Pre-BCR signaling regulation mediated by sialylated receptors recruitment as compared to BCR signaling. **(a)** CD22 dependent regulation of the B-cell receptor (BCR) signal (reviewed in^66^). CD22 forms oligomers by binding to sialylated receptors distinct from the BCR^28^. Upon specific antigen binding, conformational opening of the BCR occurs, followed by BCR activation and signaling through phosphorylation of Igα/β complex. After activation, CD22 receptors are recruited to the BCR by cytoplasmic effectors and mediate BCR signaling inhibition. **(b)** Model for pre-BCR signaling regulation. In the absence of Gal-1, “CD22-like” receptors recognizing α2,3 sialylated ligands are clustered and can trigger signaling pathways involved in cell death. When Gal-1 interacts with the pre-BCR, specific targeting of α2,3 sialylated β-galactoside ligands occurs leading to receptors recruitment. The formation of a homogeneous protein platform favors efficient pre-BCR signaling leading to cell survival. In addition, this receptor segregation will create mechanical stress provoking membrane curvature and subsequent pre-BCR internalization.

At the structural level, upon λ5-UR interaction, dynamic fluctuations throughout Gal-1 occurs with an overall increased rigidity which locks Gal-1 into a conformation prone to interact with specific glycans and exclude others. This allostery-based mechanism of regulation for Gal-1/glycan interactions leads to specific selection of α2,3 sialylated glycoconjugates at the pre-B cell surface. Allostery based phenomena have been previously shown for Gal-1 binding to synthetic inhibitor molecules designed to target Gal-1 interactions^42–44^. In addition, studies on glycan recognition by Gal-1 showed that sugar binding affected residues far from the CBS and described significant changes in the dynamics of the protein thus evidencing Gal-1 conformational plasticity and allostery upon glycan interaction^45^. These data, together with our study, emphasize the existence of intramolecular communication pathways encoded within Gal-1 structure which link glycan binding to dimer interface but also to protein interaction. Rather than a simple modification of ligand targeting, these dynamic changes result in Gal-1 conversion from exo- to endo-type lectin. To the best of our knowledge, this result is the first example of such conversion dictating the glycan preference of a Galectin. Gal-1 bound to λ5-UR is able to interact with terminal and internal galactose like the tandem-repeat Gal-8^21^. Tandem-repeat Galectins (Galectins with two CRDs linked by a flexible linker) are known to be more potent in triggering cellular responses^46^, therefore λ5-UR induced conversion might increase Gal-1 efficiency at the pre-B cell surface through the formation of higher-order multimers as seen for tandem-repeat Galectins^46^.

Beyond the pre-BCR case, our results raise a fundamental question: is there a universal mechanism of allosteric modulation of glycan binding that is conserved across all Galectins through binding to non-glycosylated protein partnersã Indeed, there are increasing evidence for direct protein interactions with Galectins^16–18,47^. We also demonstrated that CXCL4 interacts with Gal-1 and control its glycan binding activities at the base of the immunoregulatory function of galectins^47^. The pre-BCR case and these latter examples all involve the three different Galectin subfamilies (prototype, chimera and tandem-repeat)^4^. Further structural, cellular and glycomic studies of these new complexes are needed to demonstrate a universal allosteric mechanism of regulation for Galectins functions. Moreover, taking into account Galectins involvement in many crucial pathologies such as cancer^6,48^ and inflammation^49^ or infection by pathogens^7^, the discovery of this selectivity switch should also increase the potential to develop molecules modulating Gal-1 interactions and function in specific pathological situations.

## Methods

### Protein production and purification

^15^N-labeled Gal-1 has been produced and purified as previously described^50^. Protein purity was checked using SDS–polyacrylamide gel electrophoresis.

### Peptide synthesis

The λ5-UR (residue 22-45) and λ5-UR-L26A-W30A peptides were chemically synthesized and purchased from Schafer-N, Copenhagen, Denmark.

### Membrane vesicle preparation

The murine stromal cell line OP9^51^ used in this study corresponds to OP9 cells whose Gal-1 expression was decreased by 75% using a Gal-1 specific shRNA (shGAL1^52^). The cells were cultivated in MEMα-Glutamax, 20% fetal calf serum (FCS), 100 U.ml^−1^ penicillin and 100 μg.ml^−1^ streptomycin. The cells were mechanically detached from the dishes using cell scrapers to avoid the use of trypsine. The human pre-B cell line Nalm6^53^ was grown in RPMI, 10% fetal calf serum, 100 U.ml^−1^ penicillin, 100 μg.ml^−1^ streptomycin and 50 μM β-mercaptoethanol. Upon centrifugation at 450g during 5 minutes at room temperature, cells were re-suspended with PBS (Phosphate Buffer Saline) 1X (137 mM NaCl pH 7.4, 2.7 mM KCl, 4.3 mM Na_2_HPO_4_, 1.47 mM KH_2_PO_4_). Cells were vesiculated by sonication and subsequently centrifuged at 100,000g for 1h at 4°C to collect membrane vesicles. Vesicles were resuspended in 20 mM KPO_4_ pH 5.2 buffer. Protein concentration is then estimated using the Pierce™ BCA protein assay kit.

### Negative staining electronic microscopy

Experiments were performed using samples of pre-B or stromal cell vesicles placed on glow-discharged carbon-coated grid during 60 s and the excess solution was removed by touching a piece of filter paper to the side of the grid. After the adsorption of the sample on the support, the grid was washed by being placed on the surface of a drop of deionized water. This step was repeated three times before the grid was blotted dry with filter paper. Three drops of filtered (0.02µm filter) 2% uranyl formate solution were applied on the sample during 3 minutes for each drop before the excess solution was removed by blotting similarly between each drop application. The grid was then air dried. The uranyl formate being light-sensitive, the filter syringe was wrapped in aluminum foil to reduced light exposition. Images were recorded on an FEI Tecnai 200-kV electron microscope.

### PNGase F and O-glycosidase cell treatments

The cells were resuspended in 50 mM sodium phosphate buffer, centrifuged 5 minutes (450*g*) at room temperature and the supernatant was discarded. This step was repeated twice to allow the removal of the cell media growth. The PNGase F (New England Biolabs) and O-glycosidase (New England Biolabs) enzymes were added (50 000 U and 1 800 000 U, respectively) to 10 million resuspended stromal cells and incubated overnight at 37°C. After centrifugation 5 minutes at 450*g* at room temperature, the treated cells were washed twice with PBS buffer. Protein concentration is then estimated before storage at −80°C. The control of the treatment consisted in incubating 1 hour the treated and untreated cells with His-tagged Gal-1 (His-Gal-1) followed by washing steps. The binding of the His-Gal-1 to treated and untreated cells was assessed by Western Blot.

### Cell viability assays

Before treatments, cells were starved with RPMI 0,5% FCS, 100 U.ml^−1^ penicillin and 100 μg.ml^−1^ streptomycin during 1h30 at 37°C, 5% CO_2_. After two washes with cold PBS, 1.10^6^ cells were treated with 10μg/mL MAL II lectin (Eurobio, L-1260-2) alone or with 10μg/mL Galectin-1 or with 10μg/mL Galectin-1 and 100μg/mL λ5-UR peptide (Schafer-N) during 30min on ice. Then, cells were centrifugated 5 min, 450g at 4°c and the pellet was resuspended with cold PBS. Pre-B Cell viability was assessed using trypan blue dye exclusion test^54^ and the CellTiter-Glo® assay (Promega, G7570). For the CellTiter-Glo assay, CTG reagent was mixed at a 1:1 volume ratio with cells on a plastic 96-well plate according to the manufacturer’s protocol and luminescence measurements were completed using the TECAN (Männedorf, Suisse).

### NMR spectroscopy

All NMR experiments have been recorded on a 600 MHz Bruker™ AVANCE III spectrometer equipped with a TCI cryoprobe at 303 K. All samples were in 20 mM phosphate buffer at pH 5.2. All NMR experiments were recorded using 5 mm NMR tubes except experiments on ^13^C-labelled SdiLN which were recorded using 3 mm NMR tubes. All spectra were analyzed using ccpNMR Analysis software^55^, and the dynamics spectra experiments were analyzed using the Bruker™ Dynamic Center software.

### NMR titrations

The NMR titrations of Gal-1/vesicles interactions were performed using a series of 2D ^1^H, ^15^N HSQC NMR spectra recorded on a sample of purified ^15^N-labelled Gal-1 at 200 µM concentration and increasing amounts of cellular vesicles. Final titration points for pre-B and stromal cell vesicles correspond to the addition of vesicles containing 2 mg proteins. To obtain this amount of protein 100 million pre-B and 10 million stromal cells have been used. The same series were performed in the presence of 900 µM λ5-UR peptide. NMR data were analyzed using Gal-1 NMR resonance assignment previously published^56^. The chemical shift perturbations for each resonance were calculated using the equation: Δ*δ*_obs_=[Δ*δ*_HN_^2^+(Δ*δ*_N_^2^/25)]^1/2^ where Δ*δ*_HN_ and Δ*δ*_N_ are, respectively, the proton and nitrogen chemical shift variations of each residue.

### STD experiments

2D ^1^H-^13^C STD HSQC experiments were acquired with a saturation time of 2 seconds with 256 increments of 128 scans each. The STD experiments were performed using 100:1 SdiLN/Gal-1 and 100:1:3 SdiLN/Gal-1/λ5-UR molar ratio samples at 1 mM concentration for SdiLN. For 1D and 2D STD spectra the on-resonance irradiation was at 6.7 ppm and the off resonance irradiation at 20 ppm and were performed with a saturation time of 2 s.

### NMR relaxation measurements

Dynamics of ^15^N-Gal-1 free and bound to λ5-UR peptide or Anginex peptide was obtained using NMR experiments acquired at 60.81 MHz for the ^15^N frequency. The longitudinal (*T*_1_), transverse (*T*_2_), and ^1^H–^15^N-heteronuclear NOE spin relaxation times for the backbone ^15^N atoms of Gal-1 free and bound to λ5-UR peptide or Anginex peptide were collected at 303 K using the well-established NMR pulse sequences described previously^57–59^. The *T*_1_ and *T*_2_ relaxation times were measured using the following series of the delays: 10, 20, 50, 100, 200, 300, 400, 600, 800, 1000, 1500 and 2000 ms for *T*_1_ and 17.6, 35.2, 52.8, 70.4, 88.0, 105.6, 123.2, 140.8, 158.4 and 176 ms for *T*_2_. The relaxation rates R1 and R2 were calculated by fitting the decay curves to a two-parameter single exponential decay function using the Bruker software Dynamic center. ^1^H-^15^N heteronuclear NOE were measured from the HSQC and the value were calculated as the peak intensity ratio with or without amide proton saturation.

### General synthesis procedure for N-Acetylneuraminyl-α-2,3-D-Galactopyranosyl-β-1,4-2-N-Acetomido-D-glucopyranosyl-β-1,3-D-Galactopyranosyl-β-1,4-N-Acetyl-D-Glucosamine (SdiLN)

#### Materials

All non-stable isotope labeled nucleotides, nucleotide-sugars, enzymes, and chemicals were purchased from Sigma-Aldrich (St. Louis, MO), unless otherwise stated. Ni-NTA Agarose was purchased from Qiagen (Santa Clarita, CA), and Bio-Gel P2 from Bio-Rad (Hercules, CA). Glucose (^13^C_6_, 99%) was purchased from Cambridge Isotope Laboratories (Andover, MA). The enzymes used in the synthesis of SdiLN were prepared according published methods: GalE^60^, HP-39^61^, PmST1^62^. The UDP-Glucose (^13^C_6_, 99%) was prepared by previously published methods^63^.

#### D-Galactopyranosyl-β-1,4-N-Acetyl-D-Glucosamine (Supplementary Fig. 6a, compound 2)

To a reaction mixture (400 µL) containing 40 mM N-Acetyl-D-Glucosamine (*Supplementary Fig. 6a*, compound 1, 16 µmol), 48 mM UDP-Glc, 20 mM MnCl_2_ in 100 mM Sodium Cacodylate Buffer (pH 7.5), β4GalT1 (1 U) and GalE (10 U) were added and incubated overnight at 37 °C. The completed reaction, as evident by TLC, was centrifuged and the supernatant subjected to gel filtration on Bio-Gel P-2 (1×120 cm column, in 100 mM NH_4_HCO_3_ solution). Fractions containing 2 were combined and lyophilized to produce a white amorphous solid (∼ 12.8 µmol, ∼ 80% isolated yield).

#### 2-Acetomido-D-Glucopyranosyl-β-1,3-D-Galactopyranosyl-β-1,4-N-Acetyl-D-Glucosamine (Supplementary Fig. 6a, compound 3)

To a reaction mixture (640 µL) containing 10 mM of 2 (6.4 µmol), 12 mM UDP-GlcNAc, 25 mM KCl, 20 mM MgCl_2_, 1 mM DTT, in 100 mM Sodium Cacodylate Buffer (pH 7.5), HP-39 (10 U) was added and incubated overnight at 37 °C. The completed reaction, as evident by TLC, was centrifuged and the supernatant subjected to gel filtration on Bio-Gel P-2 (1×120 cm column, in 100 mM NH_4_HCO_3_ solution). Fractions containing 3 were combined and lyophilized to produce a white amorphous solid (∼ 4.5 µmol, ∼ 70% isolated yield).

#### D-Galactopyranosyl-β-1,4-2-Acetomido-D-glucopyranosyl-β-1,3-D-Galactopyranosyl-b-1,4-N-Acetyl-D-Glucosamine (Supplementary Fig. 6a, compound 4)

To a reaction mixture (100 µL) containing 40 mM of 3 (4 µmol), 48 mM UDP-Glc, 20 mM MnCl_2_ in 100 mM Sodium Cacodylate Buffer (pH 7.5), β4GalT1 (1 U) and GalE (10 U) were added and incubated overnight at 37 °C. The completed reaction, as evident by TLC, was centrifuged and the supernatant subjected to gel filtration on Bio-Gel P-2 (1×120 cm column, in 100 mM NH_4_HCO_3_ solution). Fractions containing 4 were combined and lyophilized to produce a white amorphous solid (∼ 3.2 µmol, ∼ 80% isolated yield).

#### N-Acetylneuraminyl-α-2,3-D-Galactopyranosyl-β-1,4-2-Acetomido-D-glucopyranosyl-β-1,3-D-Galactopyranosyl-β-1,4-N-Acetyl-D-Glucosamine: SdiLN (Supplementary Fig. 6a, compound 5)

To a reaction mixture (80 µL) containing 20 mM of 4 (1.6 µmol), 24 mM CMP-Neu5Ac, 20 mM MnCl_2_ in 100 mM Sodium Cacodylate Buffer (pH 7.5), PmST1 (1 U) was added and incubated overnight at 37 °C. The completed reaction, as evident by TLC, was centrifuged and the supernatant subjected to gel filtration on Bio-Gel P-2 (1×120 cm column, in 100 mM NH_4_HCO_3_ solution). Fractions containing 5 (SdiLN) were combined and lyophilized to produce a white amorphous solid (∼ 0.56 µmol, ∼ 35% isolated yield).

### SdiLN resonance assignment

^1^H and ^13^C NMR resonance assignments of SdiLN have been performed using 2D ^1^H, ^13^C HSQC, 2D ^1^H, ^13^C HSQC-TOCSY, 2D ^1^H, ^1^H-TOCSY, 2D ^1^H, ^1^H-ROESY and 2D ^1^H, ^1^H-COSY experiments. A sample of SdiLN with ^13^C-labeled Galactoses was used for recording 2D ^1^H, ^13^C HSQC and 2D ^1^H, ^13^C HSQC-TOCSY.

## Acknowledgments

This work was funded by grants from the Agence Nationale de la Recherche (ANR-16-CE11-0005-01), the Centre National de la Recherche Scientifique and the Aix-Marseille Université. The authors thank the participation of the Protein-Glycan Interaction Resource of the CFG (supporting grant R24 GM098791) and the National Center for Functional Glycomics (NCFG) at Beth Israel Deaconess Medical Center, Harvard Medical School (supporting grant P41 GM103694). This work has benefited from the facilities and expertise of the Platform for Microscopy of the Mediterranean institute of microbiology, the authors thank especially Artemis Kostas.

## Author Contributions

P.T. and L.E. designed research. P.T., B.S., A.C., O.B., Q.C., L.G.S., S.J.M. and L.E. performed research. J.R.W contributed new reagents. P.T., C.S.K. and L.E. analyzed data. L.E. wrote the manuscript.

## References

1. Varki, A. Biological roles of glycans. Glycobiology 27, 3–49 (2017).

2. Laine, R. A. Invited Commentary: A calculation of all possible oligosaccharide isomers both branched and linear yields 1.05 × 1012 structures for a reducing hexasaccharide: the Isomer Barrier to development of single-method saccharide sequencing or synthesis systems. Glycobiology 4, 759–767 (1994).

3. André, S., Kaltner, H., Manning, J. C., Murphy, P. V. & Gabius, H.-J. Lectins: Getting Familiar with Translators of the Sugar Code. Molecules 20, 1788–1823 (2015).

4. Johannes, L., Jacob, R. & Leffler, H. Galectins at a glance. Journal of Cell Science 131, jcs208884 (2018).

5. Leffler, H. Galectins structure and function--a synopsis. Results Probl Cell Differ 33, 57–83 (2001).

6. Girotti, M. R., Salatino, M., Dalotto-Moreno, T. & Rabinovich, G. A. Sweetening the hallmarks of cancer: Galectins as multifunctional mediators of tumor progression. Journal of Experimental Medicine 217, e20182041 (2019).

7. Vasta, G. R. Galectins in Host–Pathogen Interactions: Structural, Functional and Evolutionary Aspects. in Lectin in Host Defense Against Microbial Infections (ed. Hsieh, S.-L.) 169–196 (Springer, 2020). doi:10.1007/978-981-15-1580-4_7.

8. Nio-Kobayashi, J. & Itabashi, T. Galectins and Their Ligand Glycoconjugates in the Central Nervous System Under Physiological and Pathological Conditions. Frontiers in Neuroanatomy 15, 74 (2021).

9. Wan, L., Hsu, Y.-A., Wei, C.-C. & Liu, F.-T. Galectins in allergic inflammatory diseases. Molecular Aspects of Medicine 79, 100925 (2021).

10. van der Hoeven, N. W. et al. The emerging role of galectins in cardiovascular disease. Vascular Pharmacology 81, 31–41 (2016).

11. Sethi, A., Sanam, S. & Alvala, M. Non-carbohydrate strategies to inhibit lectin proteins with special emphasis on galectins. European Journal of Medicinal Chemistry 222, 113561 (2021).

12. Bertuzzi, S., Quintana, J. I., Ardá, A., Gimeno, A. & Jiménez-Barbero, J. Targeting Galectins With Glycomimetics. Frontiers in Chemistry 8, 593 (2020).

13. Laaf, D., Bojarová, P., Elling, L. & Křen, V. Galectin–Carbohydrate Interactions in Biomedicine and Biotechnology. Trends in Biotechnology 37, 402–415 (2019).

14. Brewer, C. Clusters, bundles, arrays and lattices: novel mechanisms for lectin– saccharide-mediated cellular interactions. Current Opinion in Structural Biology 12, 616–623 (2002).

15. Bonzi, J. et al. Pre-B cell receptor binding to galectin-1 modifies galectin-1/carbohydrate affinity to modulate specific galectin-1/glycan lattice interactions. Nat Commun 6, 6194 (2015).

16. Advedissian, T. et al. E-cadherin dynamics is regulated by galectin-7 at epithelial cell surface. Sci Rep 7, 17086 (2017).

17. Daley, D. et al. Dectin 1 activation on macrophages by galectin 9 promotes pancreatic carcinoma and peritumoral immune tolerance. Nat Med 23, 556–567 (2017).

18. Eckardt, V. et al. Chemokines and galectins form heterodimers to modulate inflammation. EMBO Rep 21, e47852 (2020).

19. Gauthier, L., Rossi, B., Roux, F., Termine, E. & Schiff, C. Galectin-1 is a stromal cell ligand of the pre-B cell receptor (BCR) implicated in synapse formation between pre-B and stromal cells and in pre-BCR triggering. PNAS 99, 13014–13019 (2002).

20. Nagae, M. & Yamaguchi, Y. Three-Dimensional Structural Aspects of Protein– Polysaccharide Interactions. International Journal of Molecular Sciences 15, 3768–3783 (2014).

21. Moure, M. J. et al. Selective 13C-Labels on Repeating Glycan Oligomers to Reveal Protein Binding Epitopes through NMR: Polylactosamine Binding to Galectins. Angewandte Chemie International Edition 60, 18777–18782 (2021).

22. Rossi, B., Espeli, M., Schiff, C. & Gauthier, L. Clustering of Pre-B Cell Integrins Induces Galectin-1-Dependent Pre-B Cell Receptor Relocalization and Activation. The Journal of Immunology 177, 796–803 (2006).

23. Winkler, T. H. & Mårtensson, I.-L. The Role of the Pre-B Cell Receptor in B Cell Development, Repertoire Selection, and Tolerance. Frontiers in Immunology 9, 2423 (2018).

24. Elantak, L. et al. Structural Basis for Galectin-1-dependent Pre-B Cell Receptor (Pre-BCR) Activation *. Journal of Biological Chemistry 287, 44703–44713 (2012).

25. Houzelstein, D. et al. Phylogenetic Analysis of the Vertebrate Galectin Family. Molecular Biology and Evolution 21, 1177–1187 (2004).

26. Ishima, R. & Torchia, D. A. Protein dynamics from NMR. Nat Struct Biol 7, 740–743 (2000).

27. Perdicchio, M. et al. Sialic acid-modified antigens impose tolerance via inhibition of T-cell proliferation and de novo induction of regulatory T cells. PNAS 113, 3329–3334 (2016).

28. Courtney, A. H., Puffer, E. B., Pontrello, J. K., Yang, Z.-Q. & Kiessling, L. L. Sialylated multivalent antigens engage CD22 in trans and inhibit B cell activation. PNAS 106, 2500–2505 (2009).

29. Mayer, M. & Meyer, B. Characterization of Ligand Binding by Saturation Transfer Difference NMR Spectroscopy. Angewandte Chemie International Edition 38, 1784–1788 (1999).

30. Mayer, M. & Meyer, B. Group Epitope Mapping by Saturation Transfer Difference NMR To Identify Segments of a Ligand in Direct Contact with a Protein Receptor. J. Am. Chem. Soc. 123, 6108–6117 (2001).

31. Stowell, S. R. et al. Human galectin-1 recognition of poly-N-acetyllactosamine and chimeric polysaccharides. Glycobiology 14, 157–167 (2004).

32. Di Virgilio, S., Glushka, J., Moremen, K. & Pierce, M. Enzymatic synthesis of natural and 13C enriched linear poly-N-acetyllactosamines as ligands for galectin-1. Glycobiology 9, 353–364 (1999).

33. Lajoie, P. et al. Plasma membrane domain organization regulates EGFR signaling in tumor cells. Journal of Cell Biology 179, 341–356 (2007).

34. Pace, K. E., Lee, C., Stewart, P. L. & Baum, L. G. Restricted Receptor Segregation into Membrane Microdomains Occurs on Human T Cells During Apoptosis Induced by Galectin-1. The Journal of Immunology 163, 3801–3811 (1999).

35. Bednar, K. J. et al. Human CD22 Inhibits Murine B Cell Receptor Activation in a Human CD22 Transgenic Mouse Model. The Journal of Immunology 199, 3116–3128 (2017).

36. Nguyen, J. T. et al. CD45 Modulates Galectin-1-Induced T Cell Death: Regulation by Expression of Core 2 O-Glycans. The Journal of Immunology 167, 5697–5707 (2001).

37. Sacchettini, J. C., Baum, L. G. & Brewer, C. F. Multivalent Protein™Carbohydrate Interactions. A New Paradigm for Supermolecular Assembly and Signal Transduction. Biochemistry 40, 3009–3015 (2001).

38. O’Keefe, T. L., Williams, G. T., Davies, S. L. & Neuberger, M. S. Hyperresponsive B Cells in CD22-Deficient Mice. Science (1996) doi:10.1126/science.274.5288.798.

39. Nanoscale organization and dynamics of the siglec CD22 cooperate with the cytoskeleton in restraining BCR signalling. The EMBO Journal 35, 258–280 (2016).

40. Enterina, J. R., Jung, J. & Macauley, M. S. Coordinated roles for glycans in regulating the inhibitory function of CD22 on B cells. Biomedical Journal 42, 218–232 (2019).

41. Erasmus, M. F. et al. Dynamic pre-BCR homodimers fine-tune autonomous survival signals in B cell precursor acute lymphoblastic leukemia. Science Signaling (2016) doi:10.1126/scisignal.aaf3949.

42. Dings, R. P. M. et al. Antitumor Agent Calixarene 0118 Targets Human Galectin-1 as an Allosteric Inhibitor of Carbohydrate Binding. J. Med. Chem. 55, 5121–5129 (2012).

43. Dings, R. P. M. et al. Structure-Based Optimization of Angiostatic Agent 6DBF7, an Allosteric Antagonist of Galectin-1. J Pharmacol Exp Ther 344, 589–599 (2013).

44. Dings, R. P. M., Miller, M. C., Griffin, R. J. & Mayo, K. H. Galectins as Molecular Targets for Therapeutic Intervention. International Journal of Molecular Sciences 19, 905 (2018).

45. Bertuzzi, S. et al. Unravelling the Time Scale of Conformational Plasticity and Allostery in Glycan Recognition by Human Galectin-1. Chemistry – A European Journal 26, 15643–15653 (2020).

46. Earl, L. A., Bi, S. & Baum, L. G. Galectin multimerization and lattice formation are regulated by linker region structure. Glycobiology 21, 6–12 (2011).

47. Sanjurjo, L. et al. Chemokines modulate glycan binding and the immunoregulatory activity of galectins. Commun Biol 4, 1–11 (2021).

48. Elola, M. T. et al. Galectins: Multitask signaling molecules linking fibroblast, endothelial and immune cell programs in the tumor microenvironment. Cellular Immunology 333, 34–45 (2018).

49. Morosi, L. G. et al. Control of intestinal inflammation by glycosylation-dependent lectin-driven immunoregulatory circuits. Science Advances (2021) doi:10.1126/sciadv.abf8630.

50. Pace, K. E., Hahn, H. P. & Baum, L. G. Preparation of Recombinant Human Galectin-1 and Use in T-Cell Death Assays. in Methods in Enzymology vol. 363 499–518 (Academic Press, 2003).

51. Kodama, H., Nose, M., Niida, S., Nishikawa, S. & Nishikawa, S. Involvement of the c-kit receptor in the adhesion of hematopoietic stem cells to stromal cells. Exp Hematol 22, 979–984 (1994).

52. Espeli, M., Mancini, S. J. C., Breton, C., Poirier, F. & Schiff, C. Impaired B-cell development at the pre-BII-cell stage in galectin-1–deficient mice due to inefficient pre-BII/stromal cell interactions. Blood 113, 5878–5886 (2009).

53. Hurwitz, R. et al. Characterization of a leukemic cell line of the pre-B phenotype. International Journal of Cancer 23, 174–180 (1979).

54. Strober, W. Trypan Blue Exclusion Test of Cell Viability. Current Protocols in Immunology 111, A3.B.1-A3.B.3 (2015).

55. Vranken, W. F. et al. The CCPN data model for NMR spectroscopy: development of a software pipeline. Proteins 59, 687–696 (2005).

56. Nesmelova, I. V., Pang, M., Baum, L. G. & Mayo, K. H. 1H, 13C, and 15N backbone and side-chain chemical shift assignments for the 29 kDa human galectin-1 protein dimer. Biomol NMR Assign 2, 203–205 (2008).

57. Barbato, G., Ikura, M., Kay, L. E., Pastor, R. W. & Bax, A. Backbone dynamics of calmodulin studied by 15N relaxation using inverse detected two-dimensional NMR spectroscopy: the central helix is flexible. Biochemistry 31, 5269–5278 (1992).

58. Kay, L. E., Torchia, D. A. & Bax, A. Backbone dynamics of proteins as studied by 15N inverse detected heteronuclear NMR spectroscopy: application to staphylococcal nuclease. Biochemistry 28, 8972–8979 (1989).

59. Farrow, N. A. et al. Backbone dynamics of a free and phosphopeptide-complexed Src homology 2 domain studied by 15N NMR relaxation. Biochemistry 33, 5984–6003 (1994).

60. Chen, X., Zhang, W., Wang, J., Fang, J. & Wang, P. G. Production of α-Galactosyl Epitopes via Combined Use of Two Recombinant Whole Cells Harboring UDP-Galactose 4-Epimerase and α-1,3-Galactosyltransferase. Biotechnology Progress 16, 595–599 (2000).

61. Peng, W. et al. Helicobacter pylori β1,3-N-acetylglucosaminyltransferase for versatile synthesis of type 1 and type 2 poly-LacNAcs on N-linked, O-linked and I-antigen glycans. Glycobiology 22, 1453–1464 (2012).

62. Yu, C.-C. et al. Site-Specific Immobilization of Enzymes on Magnetic Nanoparticles and Their Use in Organic Synthesis. Bioconjugate Chem. 23, 714–724 (2012).

63. Lee, H. C., Lee, S.-D., Sohng, J. K. & Liou, K. One-pot enzymatic synthesis of UDP-D-glucose from UMP and glucose-1-phosphate using an ATP regeneration system. J Biochem Mol Biol 37, 503–506 (2004).

64. López-Lucendo, M. F. et al. Growth-regulatory Human Galectin-1: Crystallographic Characterisation of the Structural Changes Induced by Single-site Mutations and their Impact on the Thermodynamics of Ligand Binding. Journal of Molecular Biology 343, 957–970 (2004).

65. Neelamegham, S. et al. Updates to the Symbol Nomenclature for Glycans guidelines. Glycobiology 29, 620–624 (2019).

66. Meyer, S. J., Linder, A. T., Brandl, C. & Nitschke, L. B Cell Siglecs–News on Signaling and Its Interplay With Ligand Binding. Frontiers in Immunology 9, (2018).

