## Supplementary figures for "Pre-B cell receptor acts as a selectivity switch for Galectin-1 at the pre-B cell surface"

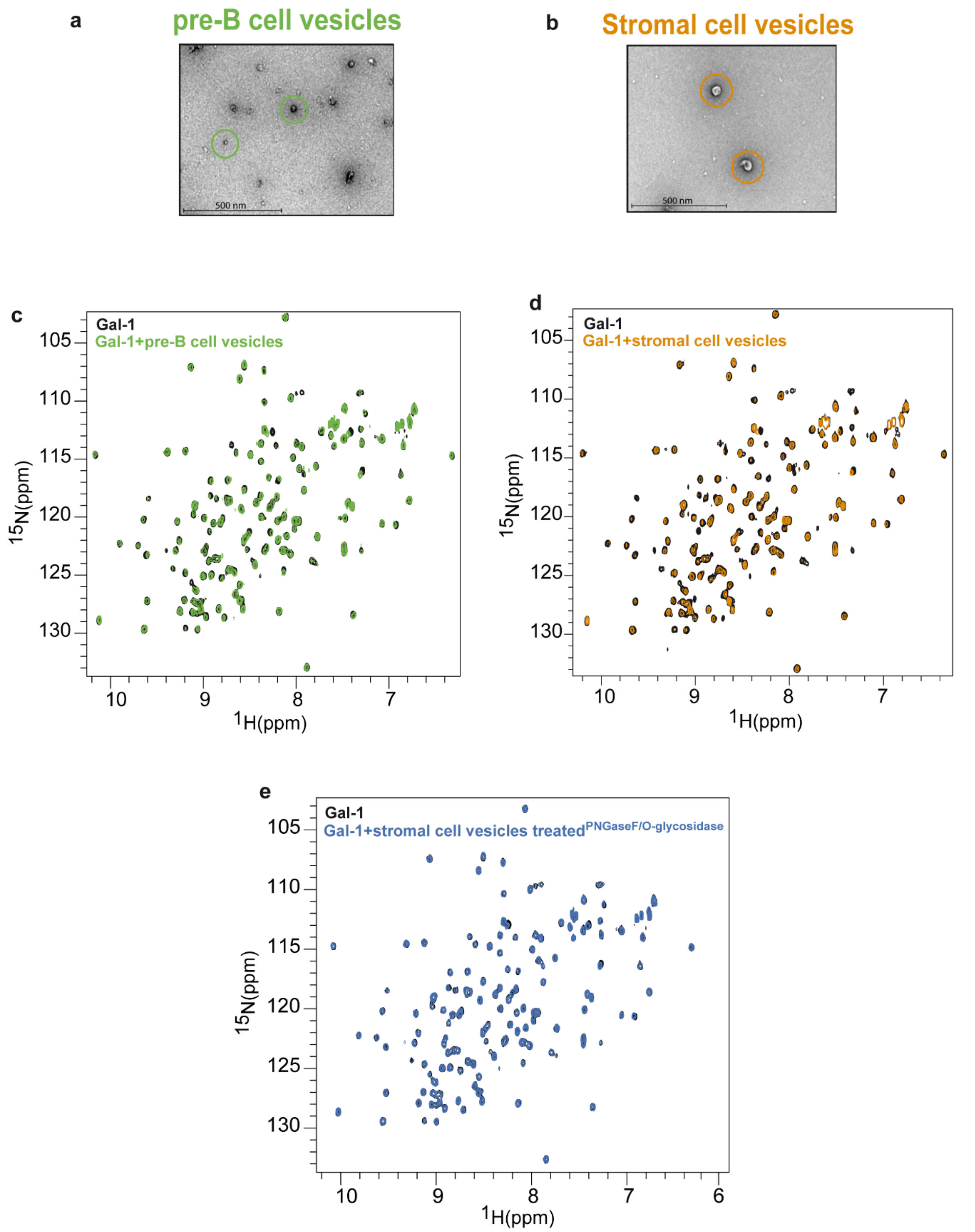


**Supplementary Figure 1. Interactions of Gal-1 with pre-B and stromal cell vesicles (a)** Negative-stain electron microscopy of the membrane vesicles extracted from pre-B and **(b)** stromal cell lines. Isolated vesicles were clearly visible and show on average a diameter of 50 nm and 25 nm for stromal and pre-B vesicles, respectively. **(c)**, **(d)** and **(e)** $^{1}H,^{15}N$ HSQC spectrum of $^{15}N$-labelled Gal-1 in the absence (black spectrum) and in the presence of membrane vesicles extracted from **(c)** 100 million pre-B cells (green spectrum), **(d)** 10 million stromal cells (orange spectrum), or **(e)** 10 million stromal cells (navy spectrum) treated with PNGase F and O-glycosidase enzymes.

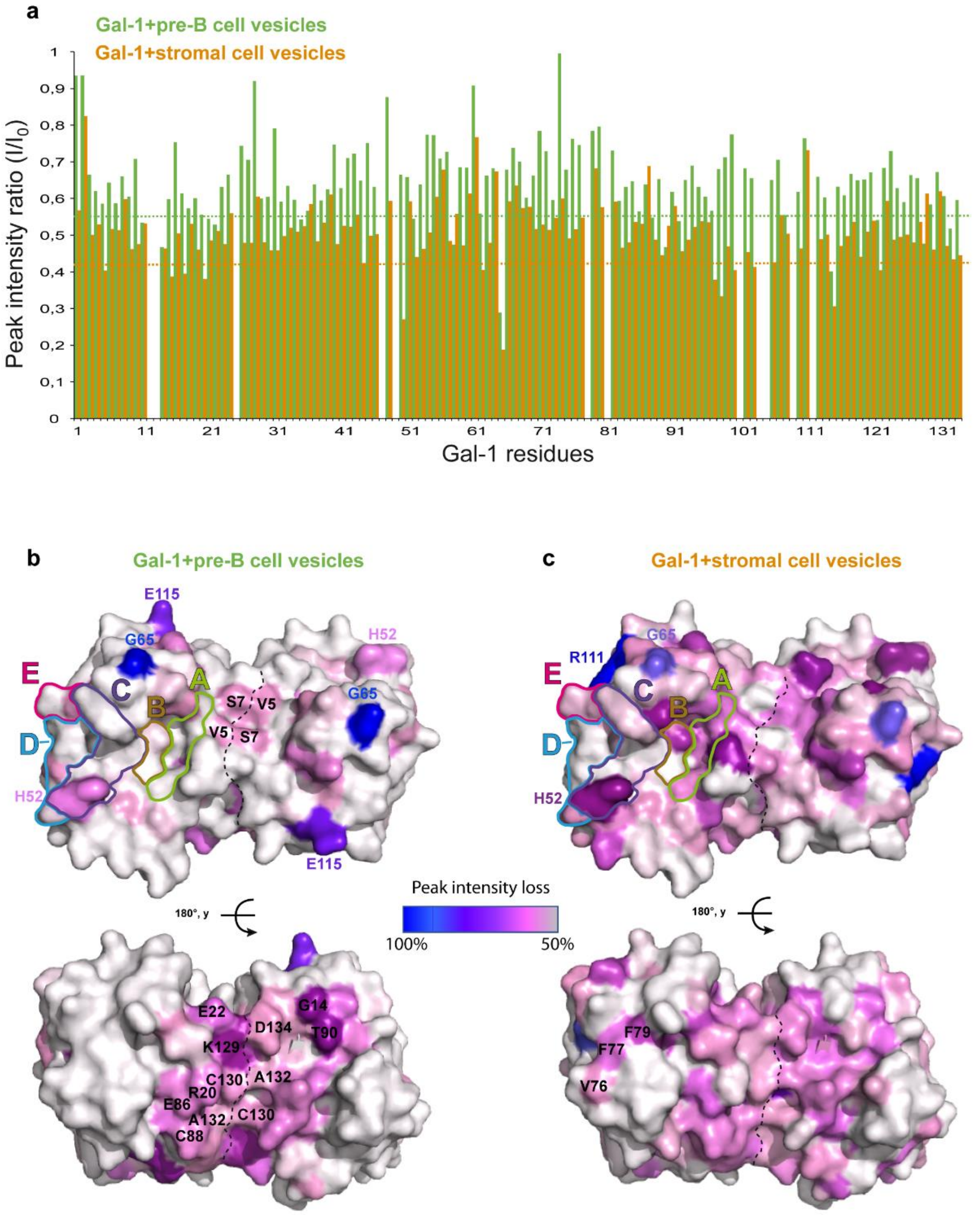


**Supplementary Figure 2. Peak intensity ratio analysis of $^{15}$N amide probes of Gal-1. (a)** Normalized peak intensity ratio analysis ($I/I_0$) of Gal-1 bound to pre-B (green bars) or stromal (orange bars) cell vesicles relative to unbound Gal-1. Dotted lines represent 1σ from the average $I/I_0$. **(b)** Two views (upper panel: CBS view, lower panel: backside view) of Gal-1 homodimer with residues affected in peak intensity analysis $I/I_0$ in the presence of pre-B cell vesicles mapped onto Gal-1. Residues are colored from blue to grey as indicated according to peak intensity loss. Dimer interface is represented with dotted lines. **(c)** Same as in **(b)** but in the presence of stromal cell vesicles.

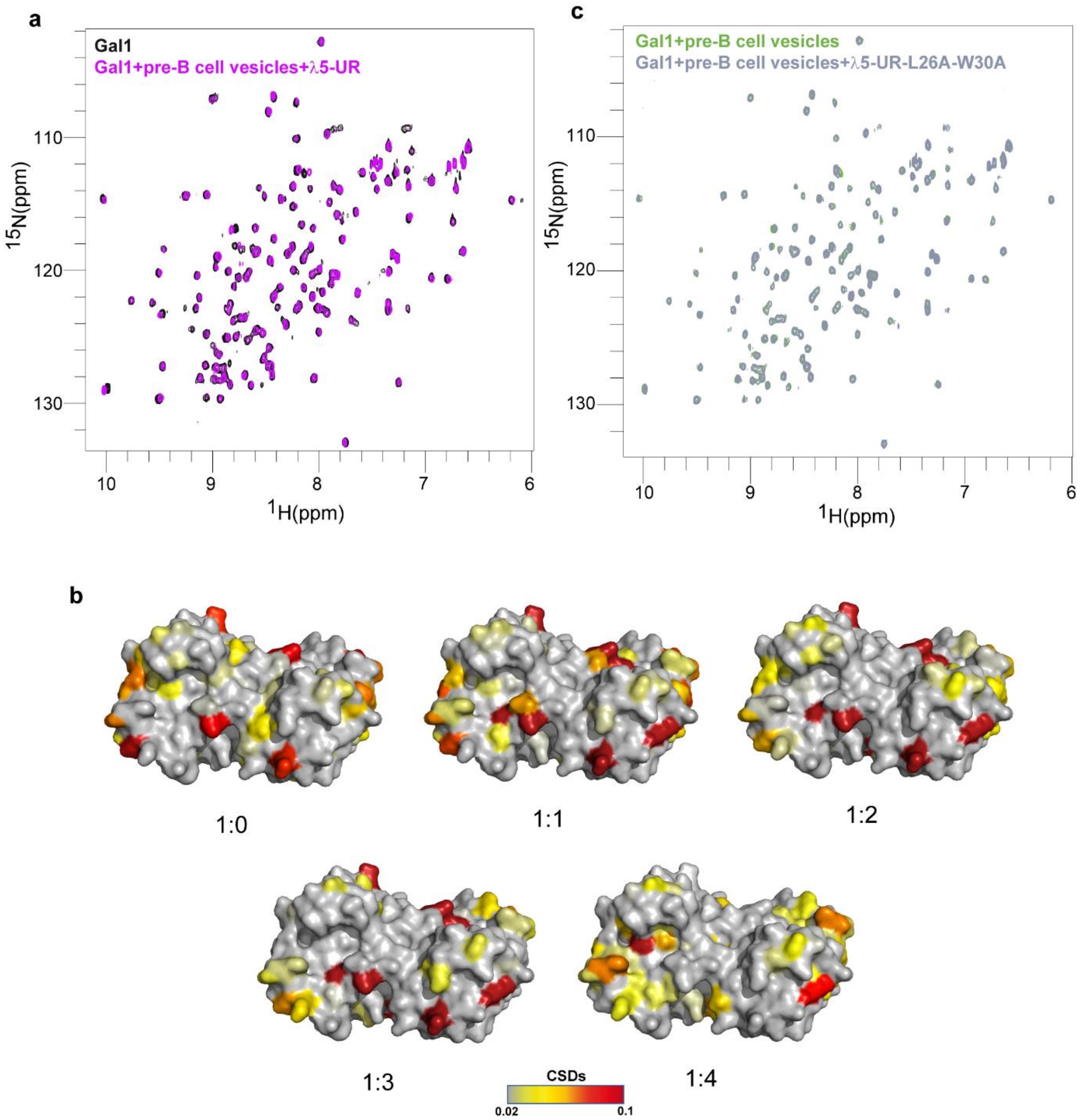


**Supplementary Figure 3. Impact of λ5-UR interaction on Gal-1 binding to pre-B cell vesicles. (a)** Overlay of $^{1}H,^{15}N$-HSQC spectrum of Gal-1 in the absence (black spectrum) and in the presence of pre-B cell vesicles and λ5-UR interacting region (magenta spectrum). **(b)** Chemical shift perturbation mapping of Gal-1 resonances in the presence of pre-B cell vesicles and increasing amount of λ5-UR. The Gal-1:λ5-UR ratio is indicated below the structures. Residues are colored from grey to red as indicated in the gradient scale. **(c)** Overlay of $^{1}H,^{15}N$-HSQC spectrum of Gal-1 in the presence of pre-B vesicles (green spectrum) and in the presence of pre-B cell vesicles and λ5-UR-L26A-W30A (grey spectrum).

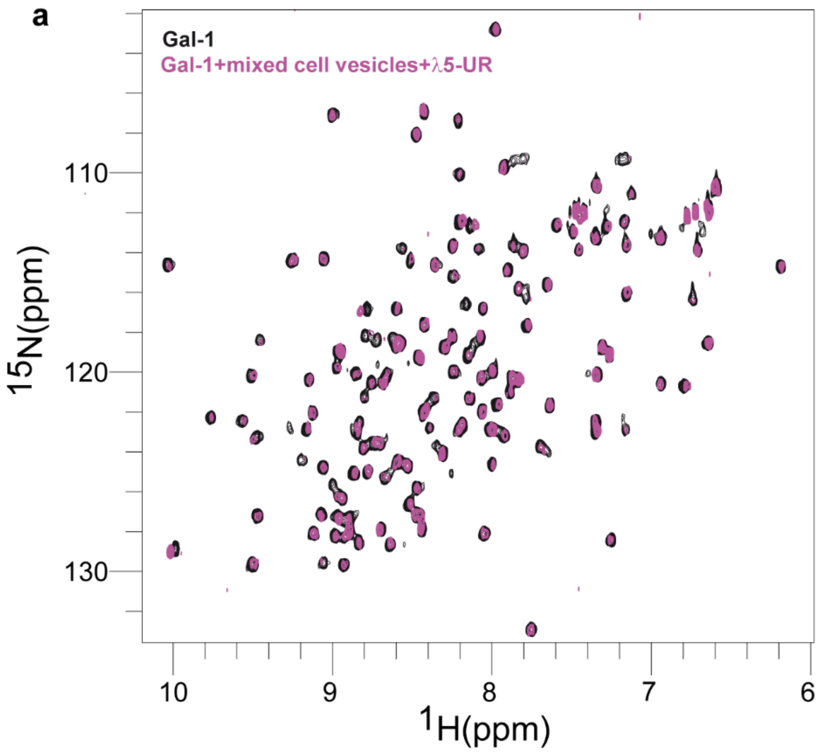


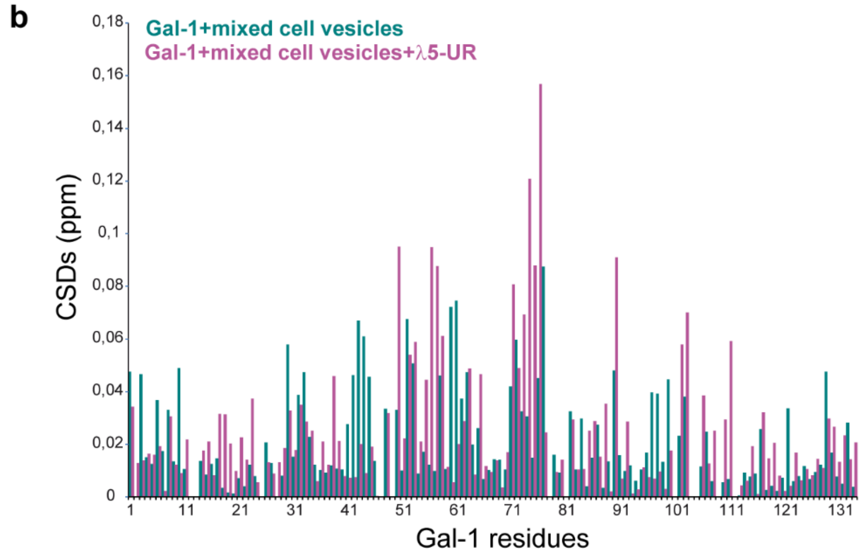


**Supplementary Figure 4. Impact of λ5-UR interaction on Gal-1 binding to mixed pre-B / stromal cell vesicles.** (a) Overlay of $^{1}$H,$^{15}$N-HSQC of $^{15}$N-labeled Gal-1 in the absence (black spectrum) and in the presence of mixed stromal and pre-B cell vesicles, and λ5-UR (purple spectrum). **(b)** Histogram plot of CSDs for Gal-1 resonances upon mixed pre-B/stromal cell vesicles interactions (teal bars), and mixed cell vesicles and λ5-UR interaction (pearly purple bars).

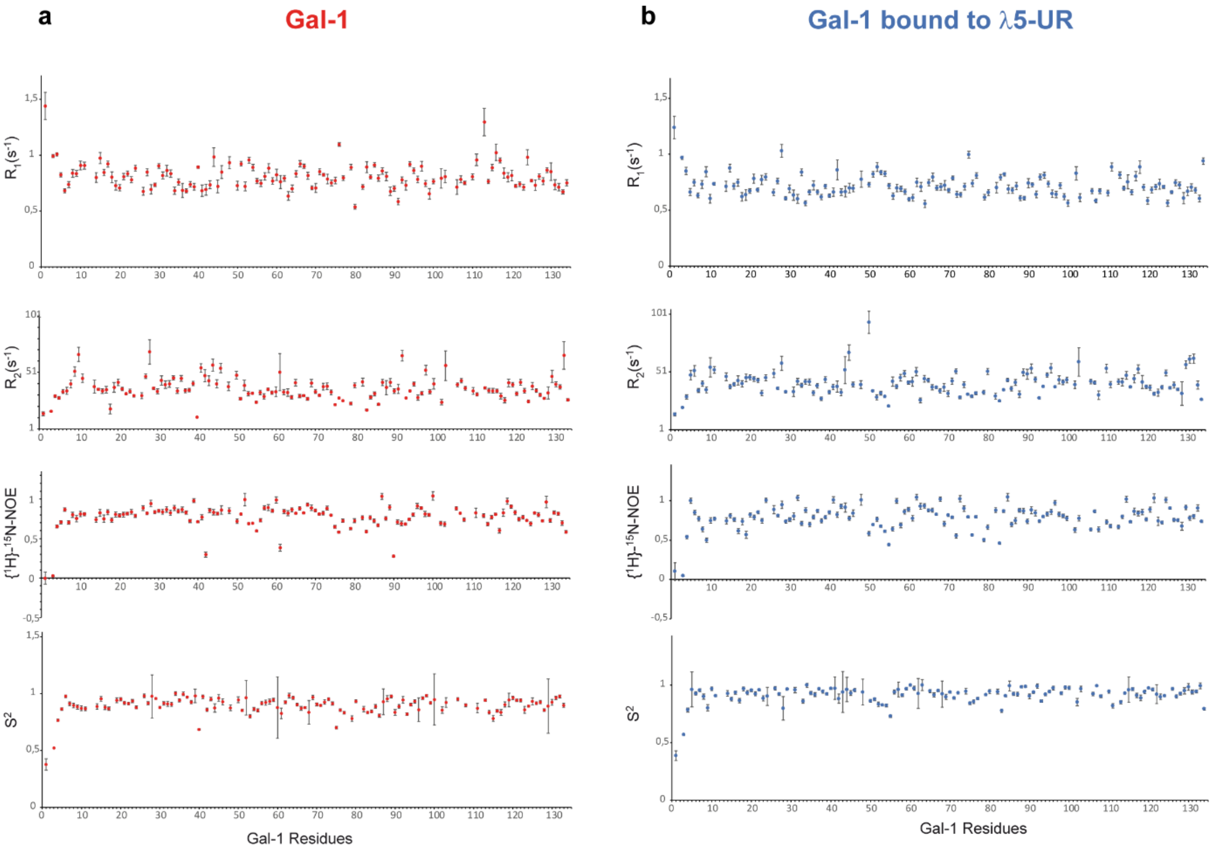


**Supplementary Figure 5:** Relaxation data for Gal-1 free and bound to λ5-UR. Relaxation rates ($R_1$, $R_2$ and $\{^1H\}$-$^{15}N$-NOE) and N-H order parameters ($S^2$) for (a) free Gal-1 and (b) λ5-UR bound Gal-1 are shown as a function of Gal-1 residue number. Data were obtained using NMR experiments acquired at 60.81 MHz for the $^{15}N$ frequency. The relaxation rates R1 and R2 were calculated by fitting the decay curves to a two-parameter single exponential decay function using the Bruker software Dynamics center. $^1H$-$^{15}N$ heteronuclear NOE were measured from the HSQC and the value were calculated as the peak intensity ratio with or without amide proton saturation. Order parameters ($S^2$) were derived using these relaxation data and the model-free approach with Dynamics center software.

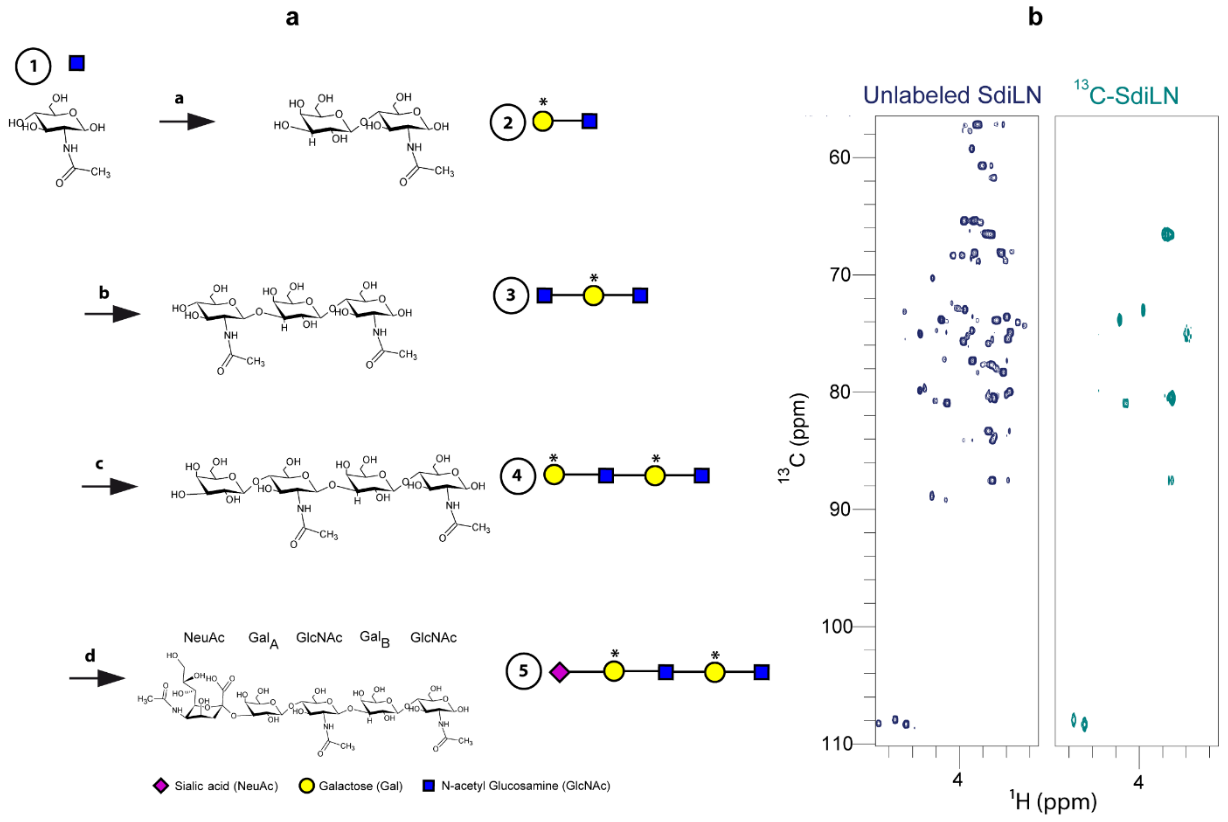


**Supplementary Figure 6. The SdiLN synthesis protocol and NMR spectra.** (**a**) General SdiLN synthesis procedure (experimental details are provided in the Methods section). Each compound structure is shown and correspond to: **1**- N-Acetyl-D-Glucosamine, **2**- D-Galactopyranosyl-β-1,4-N-Acetyl-D-Glucosamine, **3**- 2-Acetomido-D-Glucopyranosyl-β-1,3-D-Galactopyranosyl-β-1,4-N-Acetyl-D-Glucosamine, **4**- D-Galactopyranosyl-β-1,4-2-Acetomido-D-glucopyranosyl-β-1,3-D-Galactopyranosyl-b-1,4-N-Acetyl-D-Glucosamine, **5**- N-Acetylneuraminyl-α-2,3-D-Galactopyranosyl-β-1,4-2-Acetomido-D-glucopyranosyl-β-1,3-D-Galactopyranosyl-β-1,4-N-Acetyl-D-Glucosamine (SdiLN). Compound numbers are circled and their graphical representation is shown on the left (stars indicate labeled $^{13}$C-labeled Galactose). Reactions (a to d) are catalyzed using enzymes as follows: a- β4GalT1 and GalE, b- HP-39, c- β4GalT1 and GalE, d- PmST1. (**b**) Left: $^{1}$H,$^{13}$C HSQC spectrum ($^{1}$H: 3.4 - 4.7 ppm; $^{13}$C: 58 - 110 ppm) of unlabeled SdiLN. High concentration (0,5 mM) of the pentasaccharide allowed the obtention of well resolved and intense peak resonances at natural abundance. Right, same spectrum recorded on the SdiLN $^{13}$C-labeled on the six carbones of each galactose moiety.

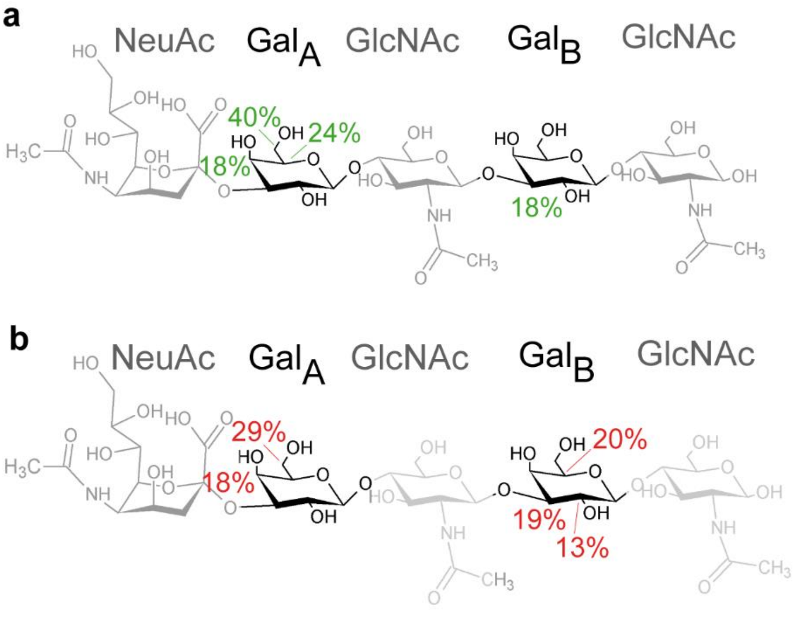


**Supplementary Figure 7. Mapping the SdiLN binding epitope using $^{13}$C-labeled SdiLN and 2D STD-NMR spectroscopy. (a)** Binding epitope of SdiLN in the presence of Gal-1. The chemical formula of the SdiLN is shown with unlabeled moieties greyed out. STD contribution has been calculated relative to the total STD signals shown in Fig. 4a, and thus expressed as a percentage of the total STD signal. These values are reported on the structure of the SdiLN. In the presence of Gal-1 alone, the strongest signal was observed for the first galactose $Gal_A$ ($Gal_A$-C4, $Gal_A$-C5, $Gal_A$-C6), suggesting that this galactose is the key component of the binding epitope. Moreover, magnetization transfer was also observed to a lesser extent for the second galactose $Gal_B$ ($Gal_B$-C3). **(b)** Same as in **(a)** but after addition of λ5-UR. Additional saturation is clearly transferred to $Gal_B$. The observed increase of the STD response suggests that λ5-UR enables enhanced Gal-1 recognition of SdiLN by allowing increased and additional intermolecular contacts with the pentasaccharide.

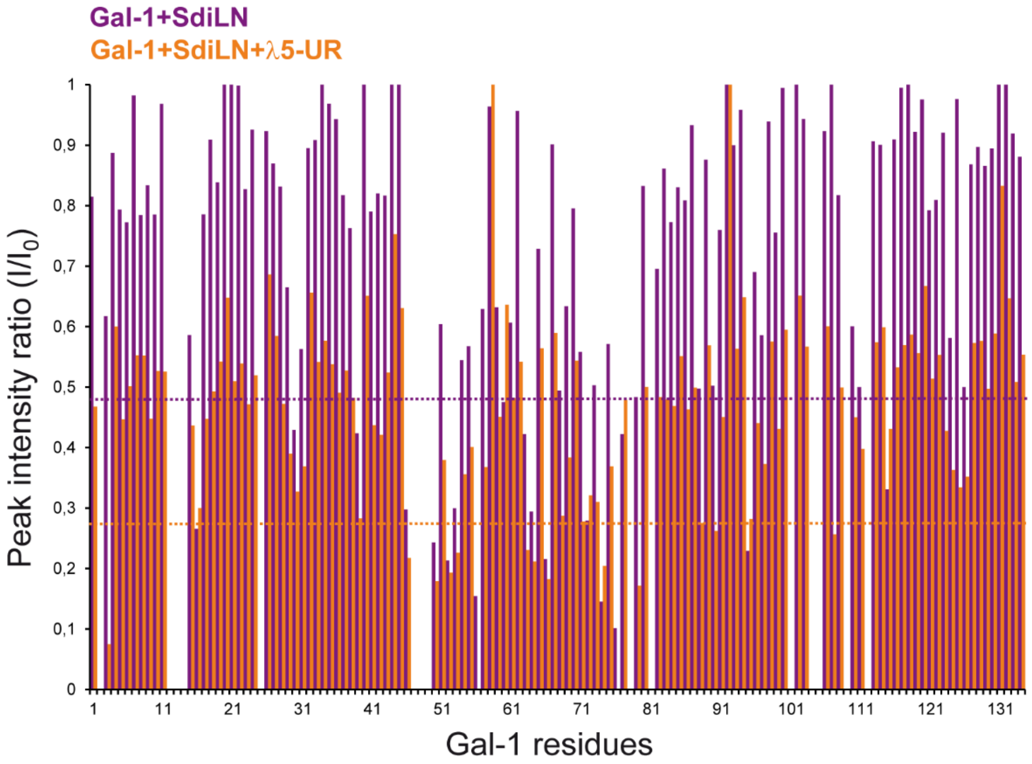


**Supplementary Figure 8. Peak intensity ratio analysis of $^{15}N$ amide probes of Gal-1 in the presence of SdiLN and λ5-UR. (a)** Normalized peak intensity ratio analysis ($I/I_0$) of Gal-1 bound to SdiLN (purple bars) or SdiLN and λ5-UR (orange bars) relative to unbound Gal-1. Dotted lines represent 1σ from the average $I/I_0$.
